# Proteomic analysis of cell cycle progression in asynchronous cultures, including mitotic subphases, using PRIMMUS

**DOI:** 10.1101/125831

**Authors:** Tony Ly, Arlene Whigham, Rosemary Clarke, Alejandro Brenes-Murillo, Brett Estes, Patricia Wadsworth, Angus I. Lamond

## Abstract

The temporal regulation of protein abundance and post-translational modifications is a key feature of cell division. Recently, we analysed gene expression and protein abundance changes during interphase under minimally perturbed conditions (Ly et al. 2014; Ly et al. 2015). Here we show that by using specific intracellular immunolabeling protocols, FACS separation of interphase and mitotic cells, including mitotic subphases, can be combined with proteomic analysis by mass spectrometry. Using this PRIMMUS (PRoteomic analysis of Intracellular iMMUnolabeled cell Subsets) approach, we now compare protein abundance and phosphorylation changes in interphase and mitotic fractions from asynchronously growing human cells. We identify a set of 115 phosphorylation sites increased during G2, which we term ‘early risers’. This set includes phosphorylation of S738 on TPX2, which we show is important for TPX2 function and mitotic progression. Further, we use PRIMMUS to provide a proteome-wide analysis of protein abundance remodeling between prophase, prometaphase and anaphase.

## Introduction

The mitotic cell division cycle is composed of four major phases, i.e., G1, S, G2 and M. The phases are defined by two major events during cell division: DNA replication (S phase) and mitosis (M phase), with intervening gap phases (G1 and G2). The cell cycle is driven by the expression of key proteins, called cyclins. Generally, cyclin expression and function is restricted to specific cell cycle phases, driving temporally ordered phosphorylation of key substrates by interacting with their kinase partners, the cyclin dependent kinases (CDKs). Temporally regulated degradation of the cyclins ensures that progression through the cell cycle is unidirectional. For example, cyclin A expression increases during S-phase, reaching a maximum in mitosis. During prometaphase, cyclin A is targeted for degradation by the anaphase promoting complex/cyclosome (APC/C), a multiprotein E3 ubiquitin ligase, thus restricting cyclin A-driven phosphorylation to S, G2 and early M-phase.

Mitosis can be further resolved into subphases (i.e. prophase, prometaphase, metaphase, anaphase, telophase & cytokinesis), which are characterised by the widespread reorganization of subcellular architecture. For example, in prophase, duplicated centrosomes separate to form the poles of the mitotic spindle. Centrosome separation is dependent on the activities of several kinases, including Cdk1 and Plk1 (Smith et al. 2011), and on the microtubule motor protein Eg5 (Sawin et al. 1992). Improperly timed centrosome separation can lead to chrosomomal instability, as shown in cyclin B2-overexpressing MEFs (Nam & van Deursen 2014). From prophase to prometaphase, nuclear envelope breakdown occurs and spindle assembly begins. During prometaphase, proper kinetochore microtubule attachments form, which depend on the interaction of each kinetochore on the sister chromatids forming stable, end on attachments to spindle microtubules. The recruitment of Eg5 to spindle microtubules is promoted by the microtubule-nucleating protein TPX2 (Eckerdt et al. n.d.; Ma et al. 2010; Ma et al. 2011). Indeed, expression of a TPX2 mutant lacking the Eg5-interaction domain leads to defects in spindle assembly and mitotic arrest (Ma et al. 2011).

Scheduled degradation of proteins is crucial for linking chromosome alignment and segregation (King et al. 1996) and is therefore an important regulatory mechanism for maintaining genome stability during cell division. Indeed, disruption of the scheduled degradation of key proteins, such as cyclins A and B, can lead to nuclear abnormalities, hyperplasia (Bortner & Rosenberg 1995) and chromosomal instability (Nam & van Deursen 2014).

A major challenge with the biochemical analysis of mitosis is that each mitotic subphase is exceedingly short, typically lasting only minutes (Sullivan & Morgan 2007), with cell-to-cell variability in dwell times. This has hampered attempts to perform a comprehensive analysis of protein abundance changes during mitotic progression due to the difficulty in obtaining highly synchronous cell populations in the various intra-mitotic stages in sufficient numbers for proteome analysis. Additionally, emerging evidence suggests that methods used to either synchronise, or arrest cells in mitosis, may induce artefacts that are not observed during an unperturbed cell cycle (Ly et al. 2015). It has also been shown that long-term spindle assembly checkpoint (SAC) activation can lead to what is termed either mitotic ‘exhaustion’, or ‘collapse’, due to sustained degradation in the presence of the active checkpoint (Balachandran et al. 2016).

Technological advances in mass spectrometry (MS)-based proteomics have enabled the large-scale detection and quantitation of proteins, including measurement of properties such as absolute abundances and subcellular localization (Larance & Lamond 2015). However, proteome measurements on either cultured cells, or tissues, reflect average values across a cell population and detailed information on biochemical state heterogeneity is obscured. A powerful approach for separating cellular subpopulations involves immunostaining combined with Fluorescence Activated Cell Sorting (FACS). CyTOF is one example combining the single cell separation power of flow cytometry with accurate mass measurement and quantitation by mass spectrometry (Bendall et al. 2011). In CyTOF, immunological reagents are conjugated with heavy metal isotopes. Cellular material is then atomized for mass analysis of individual atoms, which report on the abundance of the immunological reagent binding to cells. We note that CyTOF, while significantly increasing the number of protein antigens detected as compared with a fluorescence-based assay, does not achieve the same high depth of proteome coverage, extending to thousands of proteins, that can now be obtained by large-scale, MS-based proteomic methods (Hukelmann et al. 2016). Furthermore, apart from lower proteome coverage, CyTOF also has more limited ability to identify post-translational protein modifications and is a target-focused approach, in contrast with the unbiased analysis provided by MS-based proteomics.

Until now, studies combining FACS and MS-based proteomics have mostly involved isolating cell subpopulations based on labeling of cell surface antigens, hence avoiding cell permeabilisation and fixation (Di Palma et al. 2011),(Bonardi et al. 2013). Problems with sample bias and protein losses have been encountered in studies using MS-based proteomic methods when cells are permeabilised and fixed (Toews et al. 2008), as required for immunodetection of intracellular antigens. Indeed, to avoid intracellular immunostaining, recent work sought to identify mitosis-specific cell surface markers that can be used with live cells to isolate mitotic cell subpopulations in HeLa cells (Ozlu et al. 2015). However, as far as we are aware, neither fluorescence abundance reporter systems, nor cell surface markers, have been reported to distinguish effectively between mitotic subphases, e.g. prophase, prometaphase, metaphase & anaphase, suitable for discrimination and purification by FACS.

Previously, we reported a comprehensive proteomic dataset measuring protein and mRNA abundance variation across interphase (G1, S, and G2&M) of the cell cycle (Ly et al. 2014). To prepare cell cycle enriched cell populations, we used centrifugal elutriation (Banfalvi 2011), a method that minimizes physiological perturbation to cells and thus avoids indirect effects modulating gene expression associated with arrest procedures (Ly et al. 2015). While elutriation was effective for isolating interphase subpopulations from human leukemia cells for proteome analysis, there were limitations in resolution. In particular, mitotic cells are poorly enriched relative to G2 cells. In contrast, G2 and mitotic cells can be efficiently distinguished using intracellular immunostaining and flow cytometry (Jacobberger et al. 2008; Pozarowski & Darzynkiewicz 2004).

Building on our previous work analysing the cell cycle regulated interphase proteome (Ly et al. 2014; Ly et al. 2015), we present here a workflow for performing <u>*pr*</u>oteomics of <u>*i*</u>rntracellular i<u>*mmu*</u>nolabeled <u>*s*</u>ubpopulations of cells (PRIMMUS). Using PRIMMUS, we perform a proteome-wide analysis of changes in protein abundance and phosphorylation during interphase. Further, we perform the first proteomic characterization of distinct mitotic substages, with high enrichment efficiencies between prophase, prometaphase and anaphase in human NB4 cells. All of these proteomic data are freely available, both as raw MS files via the ProteomeXchange PRIDE repository (http://www.ebi.ac.uk/pride, TBA) and as quantified protein-level data, via the Encyclopedia of Proteome Dynamics (www.peptracker.com/EPD), a searchable, online database (Larance et al. 2013).

## Results

### Optimization of cell fixation & permeabilisation for proteome analysis

We identified three steps in intracellular immunostaining procedures that have the potential to significantly impact peptide identification by MS-based proteomics: (1) irreversibility and/or chemical modifications associated with cell fixation, (2) loss of soluble proteins during permeabalisation and (3) interference from antibody-derived peptides (Fig. 1A). Therefore, a series of experiments were performed to compare alternative fixation and permeabilisation parameters with respect to these effects.

**Figure 1.**
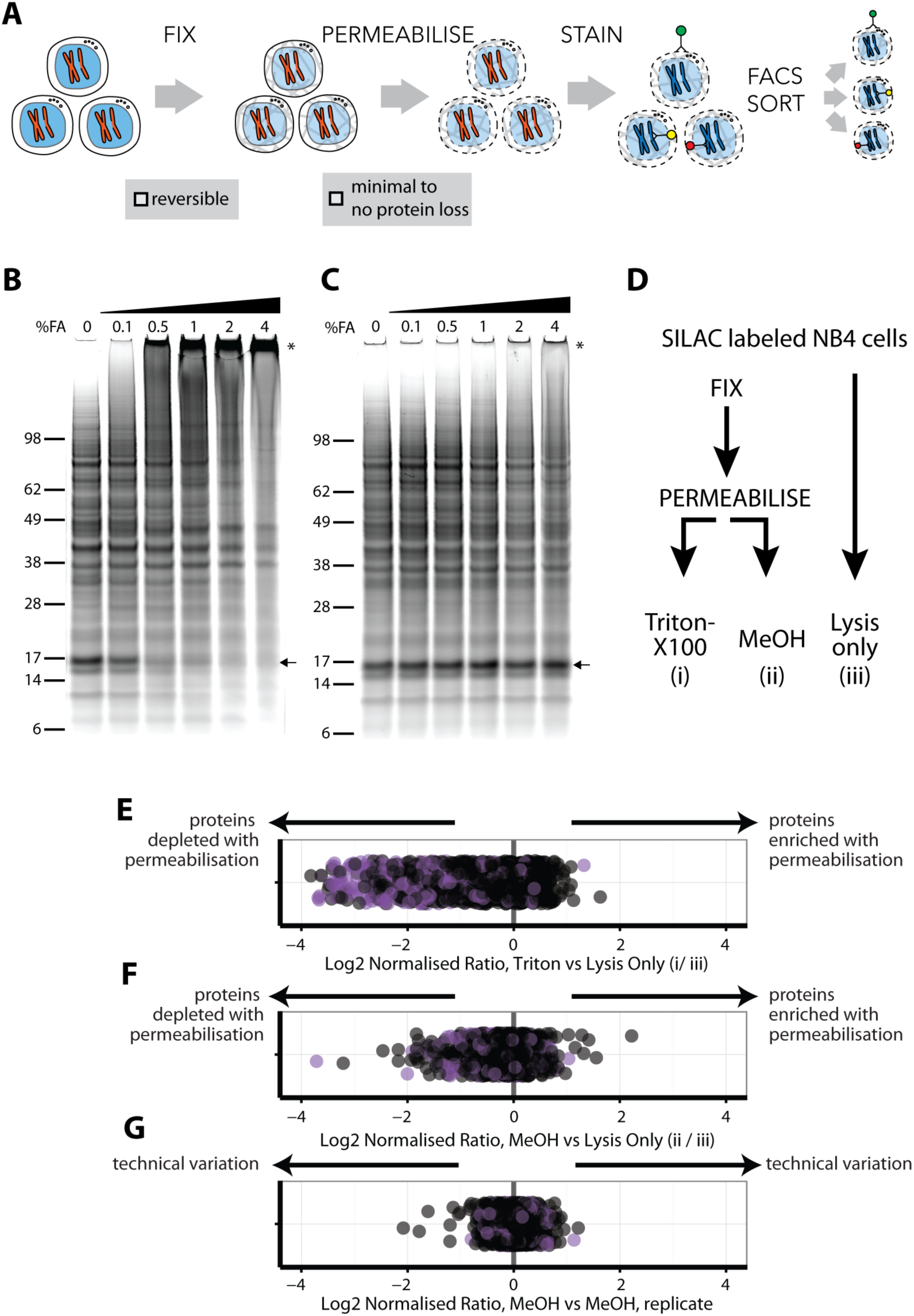
An optimised workflow for intracellular immunostaining, FACS, and MS-based proteomics. A) An abbreviated schematic of the workflow for the Proteomics of Intracellluar IMMunostained Subpopulations (PRIMMUS) approach, highlighting specific steps for optimization (fixation, permeabilisation). Lysates prepared from cells crosslinked with the indicated concentrations of formaldehyde (%v/v) in PBS (B) and then de-crosslinked with heating (C) were electrophoresed by SDS-PAGE and stained for protein using Sypro Ruby. D). SILAC labelled cells were either processed by fixation and permeabilisation, comparing 0.5% Triton X-100 (i) versus 90% methanol (ii), or with lysis only (iii). Cells were then mixed pairwise 1:1 and analysed by ‘single shot’ proteome workflows. The resulting SILAC ratios (e.g., H/L) are plotted as scatter plots for the pairwise comparisons, namely Triton X-100 vs. lysis only, methanol vs lysis only, and methanol versus methanol (technical replicate).

We chose to fix cells with formaldehyde (FA), which has been used extensively for other MS applications, such as protein-protein crosslinking (Larance et al. 2016) and crosslinked immunoprecipitations (Mohammed et al. 2016; Klockenbusch & Kast 2010). FA forms reversible crosslinks that can be broken efficiently at high temperatures. However, prior work on model peptides shows that high FA concentrations can produce irreversible chemical modifications that compromise identification by MS (Toews et al. 2008),(Sutherland et al. 2008). FA concentrations and fixation times vary significantly between common immunostaining protocols (Hutten et al. 2014). Therefore, we tested a range of FA concentrations in human myeloid leukemia NB4 cells, employing SDS-PAGE, immunoblotting and total protein gel stains to assay for crosslinking efficiency (Fig. 1B). This identified 0.5% as the minimum concentration of FA that fixes cells and produces high-MW PAGE-impermeable crosslinked products. As shown by immunoblotting, crosslinking results in ***a*** tubulin migrating at increasingly higher MW bands in a FA-dependent manner, with no monomer remaining at 4% FA (Fig. 1FS1A). A similar FA-dependent shift is observed for histone H3 (Fig. 1FS1B). These data show that while 0.1% FA is sufficient to observe crosslinked proteins, 4% FA is required to crosslink most of the total protein pool. However, high FA concentrations reduce the efficiency of reverse-crosslinking, as discussed below.

To test the efficiency of reverse-crosslinking, FA-fixed lysates were heated at 95 °C for 30 min, electrophoresed under reducing conditions and total protein visualized by SyproRuby staining (Fig. 1C). The 17-kD band that was lost in a FA concentration-dependent manner (Fig. 1B, arrow), was recovered upon crosslink reversal (Fig. 1C, arrow). However, high MW bands, indicated by an asterisk, are still observed at 4% FA after heating. Quantitation of the summed intensity within the top third of the gel indicates a ∼20% increase in intensity in 4% FA, compared with 0% FA, likely due to irreversibly crosslinked high MW protein-protein complexes. Consistent with the Sypro Ruby stain data, immunoblots show that the pools of crosslinked *α*-tubulin and histone H3 multimers, produced by treatment of cells with 4% FA, are heat stable (Fig. 1FS 1C&D).

Protein-DNA crosslinks created by FA can affect the accuracy of measuring DNA content by flow cytometry and thus may decrease the resolution with which cell cycle phases can be separated. We measured the DNA content of cells fixed with the indicated FA concentrations, permeabilised with 70% ethanol and stained with either propidium iodide (PI, left), or with 4’,6-diamidino-2-phenylindole (DAPI, right), (Fig. 1FS2). Hoechst-33342 gave inferior results (data not shown). PI staining of cells treated with 0% FA produces low coefficient-of-variation (CV) values, e.g. 5.6% for G1, which increase in a FA dose-dependent manner. Peaks for 2N and 4N DNA content (i.e., G1 and G2&M phases), coalesce at 1% and higher concentrations of FA. DAPI showed slightly higher starting CV values (7.3% for G1), but, unlike PI, was minimally affected by FA concentration. We conclude that, with these fixation conditions, 0.5% FA is the maximum for optimum use with PI, while 4% FA can be used with DAPI. In experiments where quantitation of DNA content by PI is not required to isolate a cell subpopulation of choice, then up to 2% FA can be used in combination with the intracellular staining protocol, with minimal protein loss (Fig. 1C).

Quantitative comparisons of protein levels using stable isotope labeling by amino acids in cell culture (SILAC)(Ong et al. 2002), are internally controlled, thereby minimizing the effect of technical variation after mixing on the accuracy of quantitation. We therefore used SILAC to evaluate how alternative permeabilisation reagents affect proteome recovery. For compatibility with both PI and DAPI staining and efficient crosslink reversal, cells were fixed with 0.5% FA. Three protocols were compared (Fig. 1D): (i) cells were fixed and permeabilised with 0.5% Triton X-100, (ii) cells were fixed and permeabilised with 90% methanol and (iii) cells were lysed without fixation and permeabilisation. Cells were then mixed pairwise 1:1 by cell count and processed for ‘single shot’ MS analyses (Supplementary Table 1).

The log_2_-transformed normalized ratios were analysed in scatter plots (Figs. 1E-G). In Fig. 1E, the position of each point relative to the x-axis represents the ratio between the abundance of a single protein measured in cells permeabilised with Triton X-100, (method i), and the same protein in cells processed by lysis only (method iii). Points are offset on the y-axis to minimise overplotting. Proteins detected equally in each method will show a ratio of 1, while irreversible FA-induced protein modifications and/or losses due to permeabilisation will cause lowered ratios. Known FA-induced modifications, such as methylol and Schiff base intermediates (producing +30 and +12 mass shifts, respectively), were identified after adding these modifications to the search database. However, only a small number of peptides (<1%) were found to be modified (data not shown). Proteins annotated with the subcellular localization GO terms, ‘membrane’ and/or ‘mitochondrial’, are shown in purple.

Triton X-100 is widely used in protocols for cell permeabilisation when immunostaining intracellular antigens (Hutten et al. 2014). However, we observe that many proteins are depleted in Triton-X100-treated cells (Fig. 1E). Proteins showing lower ratios after Triton-X100 permeabilisation are enriched for ‘mitochondrial’ and/or ‘membrane’ GO terms (Fisher’s exact test for enrichment: p < 1 × 10^−25^ for mitochondria and p < 1 × 10^−20^ for membrane GO annotations). In contrast, a comparison of methanol permeabilisation with direct lysis shows that most proteins vary <2-fold (Fig. 1F). However, the levels of some proteins were reduced following methanol treatment. To investigate whether this was stochastic, we compared two technical replicates of methanol-permeabilised cells using SILAC (Fig. 1G). The low variance between the replicates indicates that the effects of methanol permeabilisation on protein extraction and measurement efficiencies are systematic and reproducible. However, GO annotation analysis shows no significant enrichment for the proteins with decreased abundance after methanol permeabilisation (Fig. 1F). The reason for the selective protein loss is therefore not clear but is unlikely to significantly bias downstream proteome analyses between samples that have been similarly methanol-treated. In summary, we conclude that methanol is preferred over Triton-X100 as a permeabilisation reagent for downstream proteome analysis under these fixation conditions.

These data identify a protocol for immunostaining intracellular antigens that is compatible with efficient downstream MS-based peptide ID and quantitation and that minimizes loss of protein identifications. While the fixation and permeabilisation steps slightly reduce the overall peptide signal (Fig. 1F), these decreases are reproducible and can be accounted for in performing relative comparisons of protein abundance (Fig. 1G). We term the resulting methodology using this optimized protocol, ‘PRIMMUS’ (<u>*Pr*</u>oteomics of <u>*i*</u>ntracellular i<u>*mmu*</u>nostained cell <u>*s*</u>ubpopulations).

### PRIMMUS analysis of protein accumulation across the cell division cycle

The PRIMMUS methodology is well-suited for transforming end-point immunostaining flow cytometry assays for cell cycle analysis into a preparative procedure for global proteome characterisation. For example, G1, S and G2&M cell populations can be distinguished by DNA content alone using flow cytometry. As G2 and M phase cells have identical DNA content and similar size distributions, an additional parameter is required to separate these phases. H3S10ph, which accompanies mitotic chromatin condensation (Hendzel et al. 1997), is a specific marker for mitotic cells in many cell types and across many phyla (Hans & Dimitrov 2001). H3S10ph staining is often used as a proxy for mitotic index, particularly in flow cytometry assays (Juan et al. 1998), because only mitotic cells are H3S10ph positive. The specificity of the anti-H3S10ph antibody for mitotic cells also aids an evaluation of the potential effect of antibody IgG peptides on peptide identification.

SILAC-labeled cells were fixed, permeabilised, immunostained and sorted into four subpopulations (i.e., G1, S, G2, and M), based on both DNA content and H3S10ph staining, then processed for MS analysis, as illustrated in Fig. 2A. Four subpopulations are discernable in a representative psuedocolour scatter plot of the flow cytometric analysis, showing H3S10ph staining (y-axis), versus DNA content (x-axis) (Fig. 2B). Cells were sorted into G1, S, G2 and M fractions using the sorting ‘gates’ indicated (Fig. 2B, black boxes). The purities of G2 and M fractions were validated by co-immunostaining fractionated cells with anti-alpha tubulin antibodies and analysis by immunofluorescence (Fig. 2C). This showed that none of the cells in the G2 fraction were mitotic and >96% of cells in the M fraction were mitotic, as evaluated by chromatin condensation and microtubule organization (Fig. 2D).

**Figure 2.**
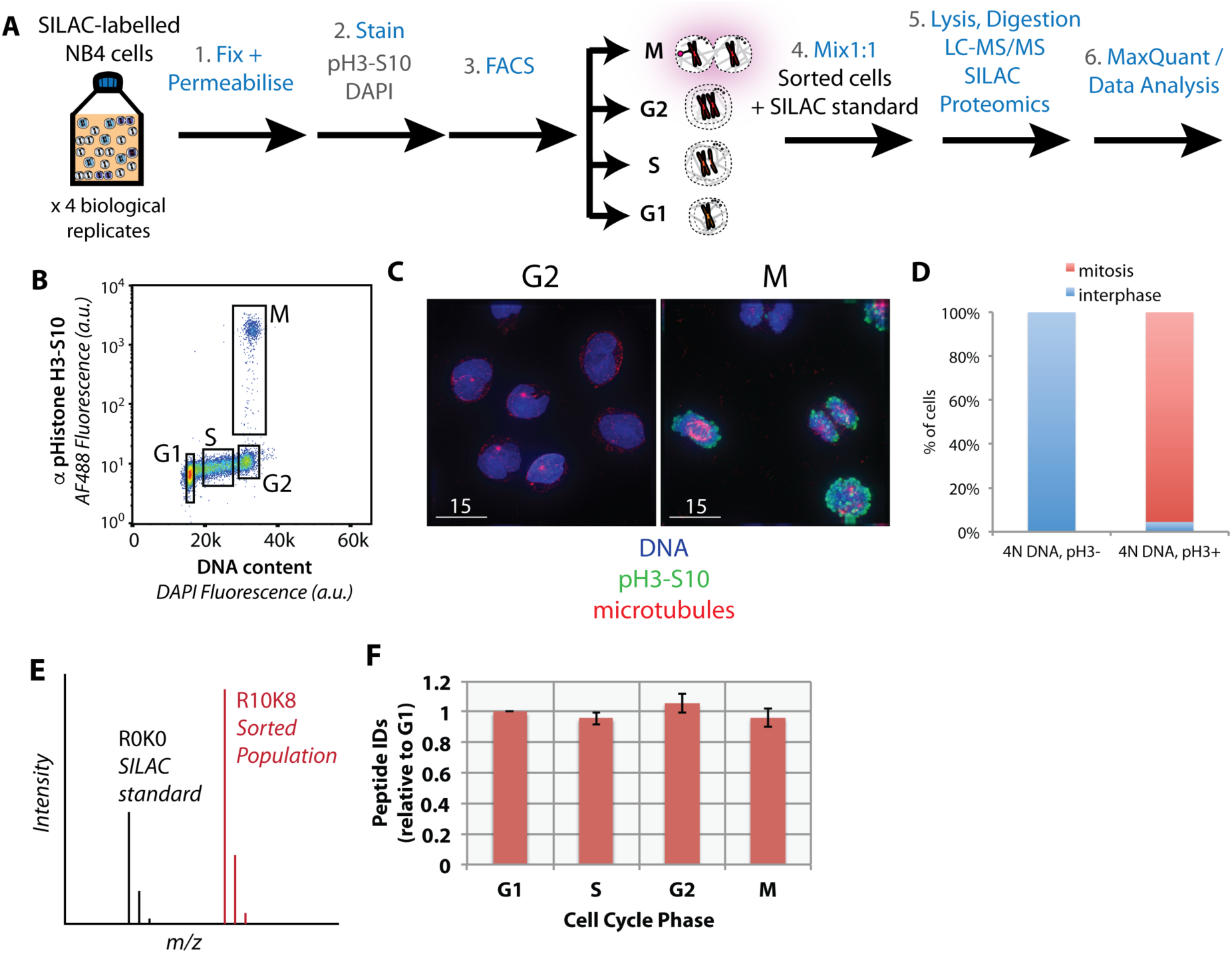
Purification of interphase and mitotic cells for PRIMMUS. Workflow for PRIMMUS of human leukemia cells into four cell cycle phase fractions (G1, S, G2, and M). Stained cells were sorted by FACS into four populations (G1, S, G2, and M) based on the gates shown on the psuedocolour plot in (B). The mitotic index of the M phase fraction was independently visualised by immunofluorescence microscopy and co-staining for microtubules (C) and quantitated (D). Fractions were then mixed by cell number 1:1 with an asynchronous SILAC-labelled standard and processed for ‘single shot’ LC-MS/MS based proteomics. The resulting measured SILAC ratios compare protein abundances in the sorted fraction versus the asynchronous standard (E). The analysis was performed with replicates (n = 4). Comparison of peptide ID rates across the sorted fractions (F).

The variation in relative protein abundance among the four sorted subpopulations of cells was determined using SILAC quantitation in single shot MS analyses (Fig. 2E). Flow sorted “heavy” cell populations were mixed with equal numbers of asynchronous “light” cells, with the signal from the “light” cells used as an internal standard to compare the four sorted populations (Fig. 2E). Four biological replicates were performed to evaluate biological variance.

In total, 32,066 peptides were identified from the four replicate experiments (raw data available at the ProteomeXchange Consortium, identifier TBA). These peptide measurements enabled quantitation of 3,696 proteins, of which 3,162 have at least two supporting peptides per protein (Supplementary Table 2). A comparable number of proteins were identified when the identical single shot MS analysis was performed on a standard complex peptide mixture prepared from directly lysed human NB4 cells (Ly et al. 2014). All cells in the asynchronous population are exposed to the H3S10ph antibody, yet only M-phase cells carry the antigen and are immunostained. Thus, the recovered M phase fractions will have IgG proteins that will be largely absent in the other fractions. Peptide ID and quantitation rates are similar across sorted populations (Fig. 1F), indicating that antibody-derived peptides present in the cell extracts from the M-phase fraction do not significantly hinder the MS-based peptide ID rate.

Because the SILAC mixing was performed using equal numbers of cells from each population, rather than equal amounts of protein, as determined either by mass, or concentration, the ratios measured here reflect the variation in average protein abundance at the respective G1, S, G2 and M phases during an unperturbed cell division cycle. Fig. 3A shows a modified violin plot (here, called a ‘neeps’ plot) of log2 ratios versus sort population (G1, S, G2 and M). The width of each ‘neep’ is proportional to the number of proteins. Black lines mark quartiles and shading indicates interquartile ranges. Compared with the internal standard (asynchronous cells), G1 cells have a median ratio of ∼0.8 and mitotic cells a median ratio of ∼1.3. Thus, an average M phase cell contains ∼1.6x the number of protein molecules present in an average G1 cell. This value is less than 2 (the approximate ratio theoretically expected between a newly divided cell and a mitotic cell), because the ‘average’ G1 cell has already spent several hours in G1 phase, during which time new protein synthesis has commenced.

**Figure 3.**
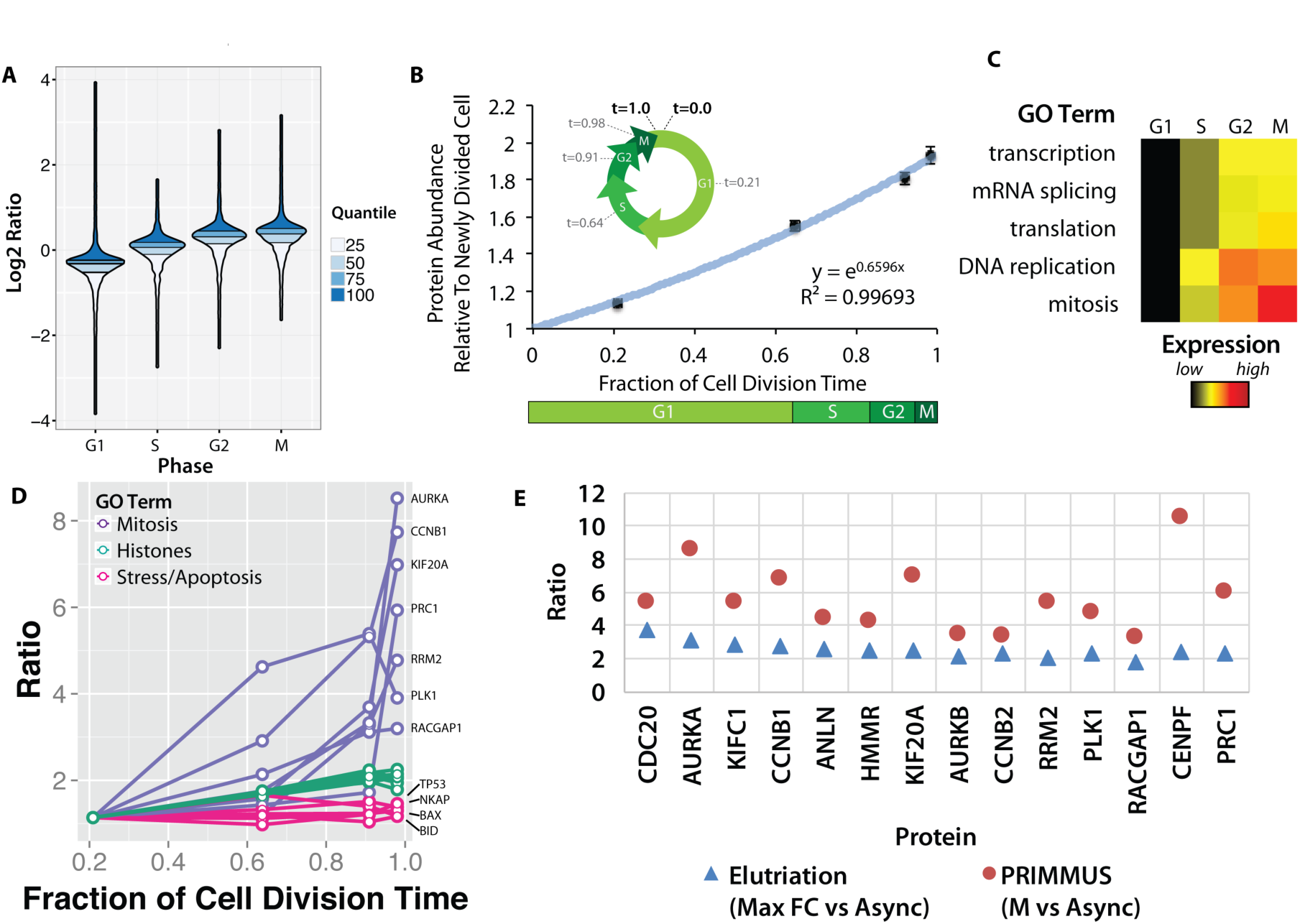
Proteomic measurement of protein accumulation across the cell division cycle. (A) A ‘neeps’ plot showing the distribution of log_2_ SILAC ratios measured in each of the G1, S, G2, and M subpopulations in one representative replicate. The width of each ‘neep’ is proportional to the density, i.e. the number of proteins. Quartiles are marked by black lines, and interquartile ranges are indicated by shading. B, inset) A schematic showing the cell division cycle and the average position during the cell division cycle for each phase collected, where a newly divided cell is defined as t = 0, and cell division (cytokinesis) is defined as t = 1. B, graph) Regression analysis was performed to produce a best-fit line in the form of an exponential growth model (i.e., y = e^mx^). (C) Ratios of proteins belonging to each of the indicated GO terms were averaged (mean) and visualized using a heatmap. (D) A plot of ratios of individual proteins associated with mitosis, chromatin, and the stress response versus cell cycle stage. E) A comparison of the ratio of G2&M vs. asynchronous measured in the elutriation vs. PRIMMUS datasets.

Using the average doubling time for NB4 cells (24 hours) and the frequency of cells in each cell cycle phase, as measured by flow cytometry, we use ergodic analysis (Pozarowski & Darzynkiewicz 2004; Wheeler 2015; Kafri et al. 2013) to estimate the average time post-division for each subset of cells and express them as a fraction of the total cell division time, i.e. t = 0 (newly divided), t = 0.21 (G1), t = 0.65 (S), t = 0.92 (G2), t = 0.98 (M) and t = 1.0 (cell division). This representation of the relative cell cycle division time (Fig. 3B, inset), allows protein abundance data to be plotted on a numerical time axis and provides a quantitative measurement of protein accumulation across the cell cycle.

Global changes in relative protein abundance at different cell cycle stages were estimated by taking the mean log2 SILAC measurement across all proteins. Mean protein measurements were calculated for each cell cycle phase, normalized to G1 and plotted as a function of cell division time, with error bars indicating the standard error of the mean (Fig. 3B). Bulk protein abundance accumulation across the cell cycle follows an exponential growth curve (r^2^ = 0.99) (Fig. 3B). Bulk protein abundance is measured to increase ∼1.9-fold between a newly divided cell and a cell about to enter the next mitosis phase. These findings are consistent with results from experiments performed decades ago, using radioactive pulse-chase experiments to analyse bulk protein accumulation (Scharff & Robbins 1965),(Rønning et al. 1979). However, using PRIMMUS, the relative protein accumulation across the cell cycle is measured here not on bulk protein alone, but rather on a per-protein basis for thousands of individual proteins.

We exploited the per-protein resolution of our data to examine whether different protein classes accumulate at different rates during the cell cycle. Fig. 3C shows a heatmap summarising an analysis of the mean normalized ratio for proteins with GO annotations for major cellular processes, i.e., transcription, mRNA splicing, translation, DNA replication and mitosis (Fig. 3C). As expected, proteins involved in DNA replication increase more on average during S phase than proteins involved in basal gene expression (transcription, mRNA splicing, translation).

We show individual profiles for selected proteins in Fig. 3D. Aurora kinase A, which regulates spindle assembly in mitosis (Floyd et al. 2008), increases in abundance by more than 8-fold between G1 and M phase (Fig. 3D). Core histones, however, show a profile that resembles bulk protein accumulation and approximately double in abundance across the cell cycle, consistent with the parallel doubling in DNA content. However, while bulk protein continues to accumulate between G2 and M (Fig. 3D), histone abundance remains relatively flat, consistent with histone protein synthesis being markedly down-regulated once S phase is completed (Fig. 3D). Interestingly, in contrast with bulk cellular proteins, we note that proteins involved in stress responses and/or apoptosis, such as p53, NKAP (a member of the NF-kappa B pathway) and the apoptotic regulators BAX and BID, remain relatively constant on a per-cell basis across these four cell subpopulations.

Using FACS, each cell cycle phase is enriched with higher purity as compared with centrifugal elutriation. We therefore analysed whether this results in the PRIMMUS method providing increased sensitivity for detecting relative changes in protein abundance between cell cycle phases. Fig. 3E shows a comparison between datasets obtained using either PRIMMUS (red circles), or elutrition (blue triangles), for 14 proteins that all peak in abundance in G2/M relative to G1 phases (i.e., CDC20, AURKA, KIFC1, CCNB1, ANLN, HMMR, KIF20A, AURKB, CCNB2, RRM2, PLK1, RACGAP1, CENPF & PRC1). All these 14 proteins show higher ratios in the PRIMMUS dataset. We conclude that the higher enrichment purities for cell subsets obtained by FACS results in PRIMMUS providing a sensitivity advantage over centrifugal elutriation.

### PRIMMUS analysis of protein phosphorylation across interphase and mitosis

We next investigated protein phosphorylation changes during interphase and mitosis using PRIMMUS. Using the identical sort strategy as above, we separated asynchronous NB4 cells into G1, S, G2 and M phase fractions by FACS (Fig. 4A). The experiment was performed in biological duplicate. The cell fractions were then processed for phosphopeptide analysis and TMT-based quantitation (8-plex, 2 biological replicates x 4 cell cycle phase fractions). Enriched phosphopeptides with no further peptide fractionation were detected and quantitated using a single shot LC-MS/MS analysis. In total, 4,500 phosphorylation sites were identified on 1,558 proteins. Most phosphorylation sites were phosphoserines (83.2%), with smaller frequencies for phosphothreonines (15.8%) and phosphotyrosine (1.0%) (Fig. 4B). Over 60% of the phosphorylation sites matched the proline-directed kinase motif (S/T followed by a proline (Fig. 4C). CDKs are one of several kinases, e.g. MAP kinases, which can phosphorylate the [S/T]P motif (as reviewed in (Amanchy et al. 2007)). The phosphoproteins detected are enriched in proteins with functions in cell division and DNA repair/stress response, in addition to ‘housekeeping’ proteins involved in gene expression (Fig. 4D). The phosphorylation site abundance profiles for the duplicate measurements were similar (Fig. 4E). As discussed in more detail below, many of the extreme differences in phosphorylation are observed in the mitotic fraction. Pearson correlations calculated for individual phosphorylation sites showed high correlation (Fig. 4F). The maximum fold changes measured for individual phosphorylation sites were also highly correlated between the two replicates, showing that the quantitation was reproducible (Fig. 4G).

**Figure 4.**
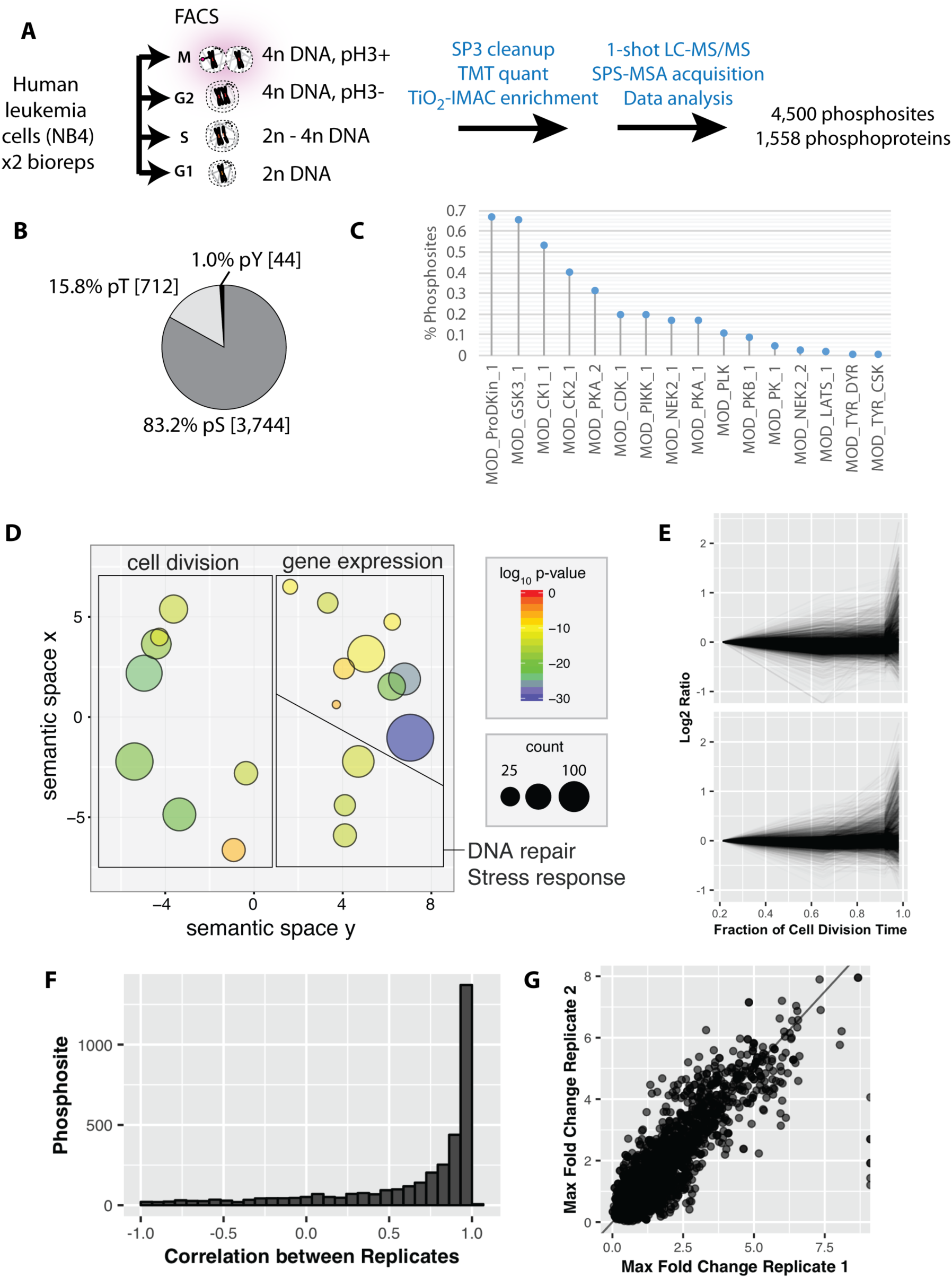
Reproducible analysis of phosphorylation changes across the cell division cycle. A) Workflow for phosphorylation analysis of FACS separated cell cycle states. B) Distribution of phosphorylated residues detected. C) Percentage of phosphorylation site sequences matching reported consensus substrate phosphorylation motifs (www.elm.eu.org, REF). D) Gene ontology enrichment analysis of the detected phosphoproteins (ReviGO, REF). E) Phosphorylation profiles across the four FACS separated fractions for replicate 1 and 2. F) A histogram of correlation coefficients calculated for individual phosphorylation sites between the two replicates. G) A comparison of measured maximum fold change between replicates.

The identified phosphorylation sites were then clustered using k-means into six groups (Fig. 5A). Cluster 1 contains phosphorylation sites that rise significantly during mitosis. Interestingly, many of these sites also show an increase in the G2 fraction (Fig. 5A, arrow). Clusters that peak in mitosis (1, 2, 3 and 5) represent 34% of the phosphorylations quantitated (Fig. 5B). Interestingly, this number is significantly smaller than reported previously in a study measuring phosphorylation dynamics in HeLa cells synchronized using nocodazole arrest and arrest-release protocols (Olsen et al. 2010). A comparison of the ratios measured in the PRIMMUS dataset with this previous analysis shows significant differences. We note that phosphorylation sites specifically upregulated in this PRIMMUS dataset are on proteins that are enriched for the gene ontology annotations ‘cell cycle’ and ‘mitosis’ (Fig. 5C, purple). In contrast, phosphorylation sites specifically upregulated in the previous HeLa cell arrest-release dataset (which are not changing in this PRIMMUS dataset), are instead on proteins showing significant enrichment for the gene ontology annotation ‘RNA splicing function’ and show no enrichment for either ‘mitosis’, or ‘cell cycle’ (Fig. 5C, cyan). These differential enrichments in specific cellular functions for the proteins identified with changing levels of phosphorylation across the cell cycle suggest underlying physiological differences between the cells used in these respective studies (see also Discussion).

**Figure 5.**
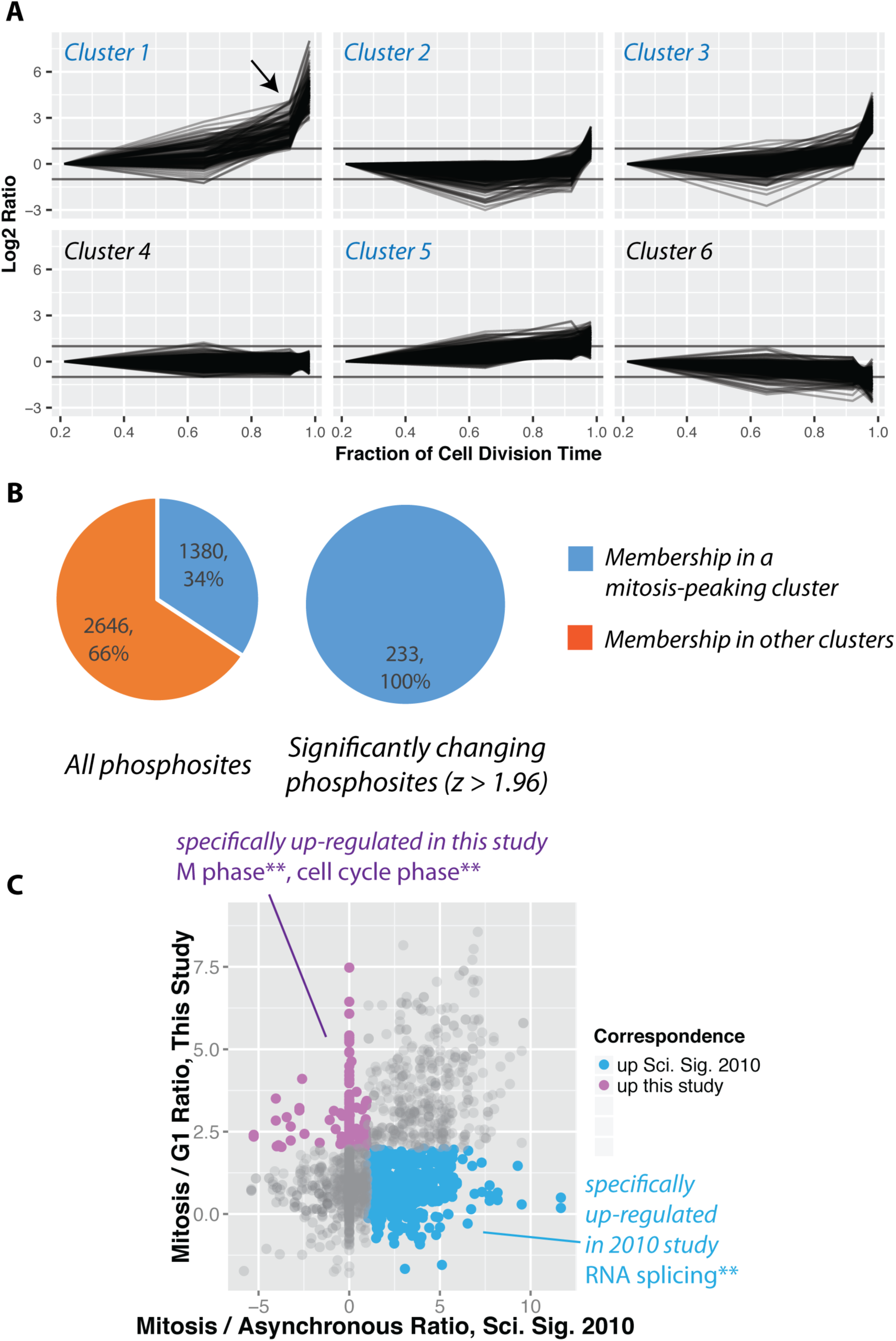
Increased global mitotic phosphorylation dominated by a subset of highly phosphorylated proteins. A) K-means clustering of the phosphorylation profiles. B) Distribution of mitosis-peaking phosphorylation sites, either in the entire dataset (left), or significantly changing phosphorylation sites (right). C) A comparison of phosphorylation site ratios measured in this dataset and a previous analysis of mitotic phosphorylation in human cells.

### Identification of a set of ‘early rising’ phosphorylation sites

While most of the significantly changing phosphorylation sites peak in mitosis, a subset, consisting of 115 sites, also show significantly increased phosphorylation in the G2 phase enriched fraction. We have termed these 115 phosphorylation sites, ‘early risers’. The high enrichment efficiency of FACS (c.f. Fig. 2C) renders it very unlikely that the increased G2 ratios measured originate from contamination from H3S10ph-positive mitotic cells in the G2-enriched fraction. Indeed, further analysis shows that early risers share additional functional similarities. Thus, early rising phosphorylation sites are situated on proteins highly enriched in nuclear, nuclear envelope and chromatin localisations (Fig. 6A). A STRING network analysis identifies several functional categories of early rising proteins, including DNA replication, cytoskeleton remodelers and spindle/kinetochore components, chromatin factors and remodelers, nuclear envelope proteins, transcription factors, nucleolar proteins and mRNA capping proteins. Proteins with the highest ranked ratios are either involved in DNA replication, or associated with the nuclear envelope (Supplementary Table 2). Motif analysis shows that early risers are enriched in the optimal CDK consensus motif of a serine/threonine followed by a proline and a downstream basic residue (Figure 6C). Consistent with a high CDK phosphorylation propensity, early risers on average have higher phosphorylation ratios during M-phase compared to ‘late risers’, (i.e. late risers being phosphorylation sites that peak in mitosis and do not increase in G2).

**Figure 6.**
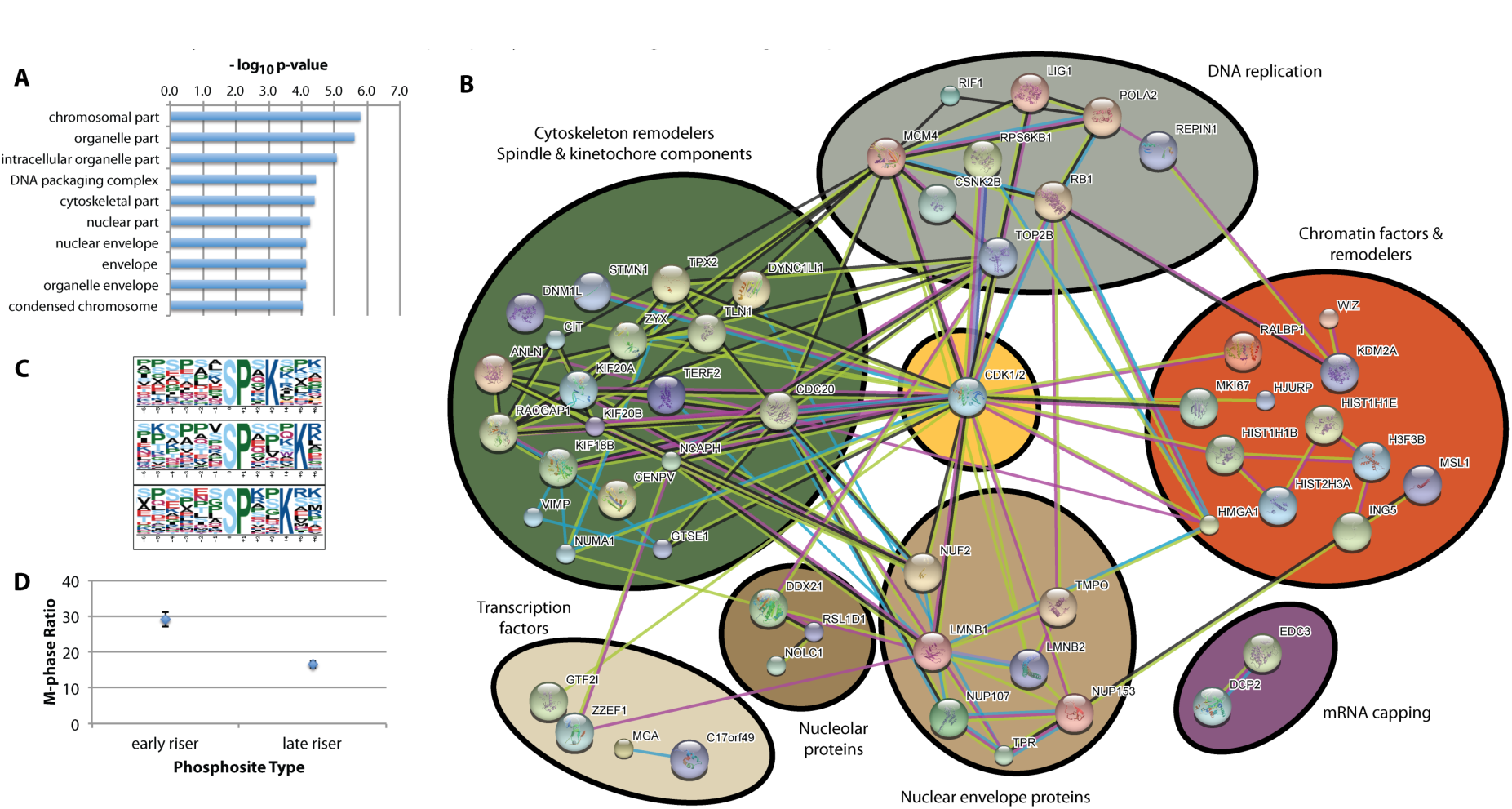
Identification of ‘early risers’, a subset of mitotic phosphorylations that begin increasing in G2 phase. A) Gene ontology enrichment analysis of early rising phosphorylation sites. B) A STRING network analysis of early rising phosphoproteins. Nodes with one or more connections are shown. C) Enriched sequence motifs among early rising phosphorylation sites (Motif-X). D) Comparing the M-phase ratio between ‘early rising’ and ‘late rising’ phosphorylation sites. Error bars show s.e.m.

One of the early risers identified was on the protein TPX2. Two TPX2 phosphorylation sites are quantitated in this dataset, i.e., S185 and S738. Both sites have surrounding sequences matching the consensus CDK phosphorylation motif (SPEK and SPK, respectively) and show increased phosphorylation in the G2-phase enriched fraction, compared with the total levels of unmodified TPX2 protein. However, TPX2 S738 is an early riser site that is phosphorylated to a greater extent in the G2 phase fraction and this difference is further increased in the M-phase fraction. We therefore decided to explore whether there was any functional relevance for phosphorylation of TPX2 at S738 in mammalian cells by analysis of phosphodefective and phosphomimetic mutants.

### Expression of S738A TPX2 mutant fails to rescue TPX2-depleted cells

TPX2 is known to have several functions during mitosis (Neumayer et al. 2014). The N-terminal domain has been shown to be important for interacting with Aurora A kinase (Bayliss et al. 2003) and this interaction facilitates microtubule nucleation at chromosomes (as reviewed in (Gruss & Vernos 2004)). The C-terminal domain has been shown to be important for interaction with Eg5. Cells expressing a C-terminal deletion mutant lacking the Eg5-interaction domain (TPX2-710) shows mislocalisation of spindle microtubule-associated Eg5 and disorganized multipolar spindles (Ma et al. 2010; Ma et al. 2011). Interestingly, the early rising phosphorylation site on TPX2, serine 738 (S738), is situated in the Eg5-interaction domain.

To assess the function of S738 phosphorylation, endogenous TPX2 was depleted in porcine LLC-Pk1 cells using siRNA and ectopic, GFP-tagged TPX2 was expressed from a rescue construct, comparing the phenotypes of either the wild-type (WT) sequence, a TPX2 phosphomimetic (S738D) mutant, or a TPX2 phosphodefective (S738A) mutant. Consistent with previous observations (Ma et al. 2011), depletion of TPX2 without rescue leads to an increase in short, collapsed spindles (Fig. 7A, right). Replacement with GFP-TPX2 WT increases the frequency of ‘normal’ bipolar spindles (Fig. 7A & 7B top). Replacement expression of the phosphomimetic TPX2 S738D mutant also rescues the spindle defect (Fig. 7A & 7B bottom), in a manner similar to replacement expression of TPX2 WT. In contrast, expression of the phosphodefective TPX2 S738A mutant results in an increase in multipolar spindles (Fig. 7A & 7B middle), a spindle defect similarly observed in cells expressing TPX2-710, a mutant missing the C-terminal Eg5-interaction domain (Ma et al. 2011).

**Figure 7.**
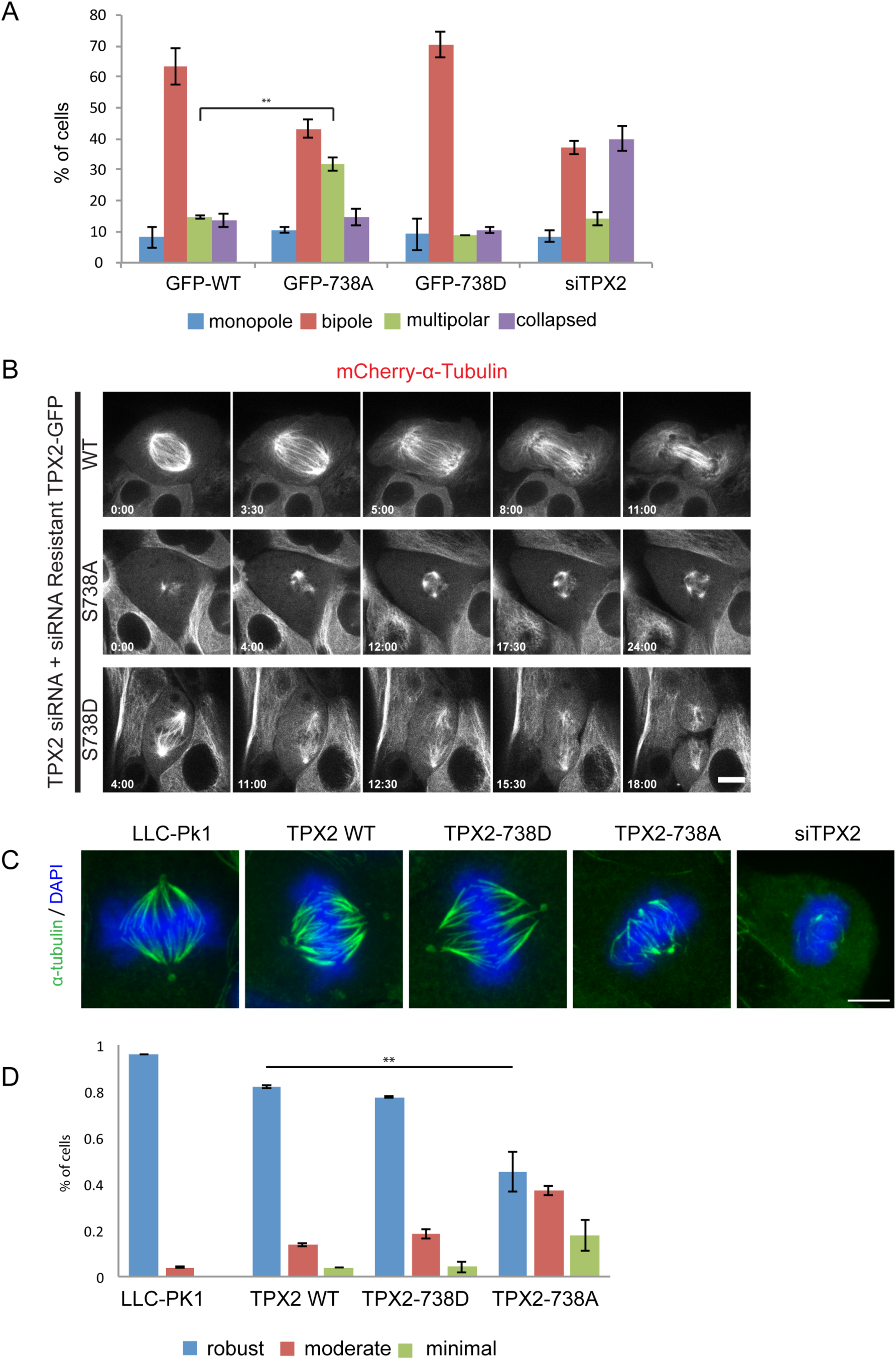
The phosphodefective S738A mutant of TPX2 fails to rescue spindle defects induced by TPX2 depletion. A) Percent of mitotic phenotypes in LLC-Pk1 cells transfected with siRNA targeting endogenous TPX2 alone or co-transfected with an siRNA resistant TPX2 construct (WT, S738A, S738D). Monopole (blue), bipole (red), muitipole (green), collapsed (purple). B) Time-lapse imaging of LLC-Pk1 cells expressing mCherry- *α*-tubulin and co-transfected with TPX2 siRNA and siRNA resistant TPX2 constructs (WT, S738A and S738D). C) Cold stability assay. Immunofluorescence staining for MTs and DNA in parental LLC-Pk1, and cells transfected with siRNA targeting TPX2 alone or co-transfected with an siRNA resistant TPX2 construct encoding wilt type (WT) TPX2, TPX2 S738A, TPX2 S738D. Maximum intensity projections of Z stacks are shown. D) Quantification of the level of cold-stable kinetochore fiber bundles shown in (C). ** P = 0.001. Scale bars in C = 5 *μ* m. Error Bars = St Dev. For D, N = 168, 151, 190, and 214 respectively.

Another closely related phenotype associated with replacement of endogenous TPX2 with TPX2-710 is that few, if any, kinetochore fibres are cold stable, indicative of either weak, or improper, kinetochore-microtubule interactions (Ma et al. 2011). We investigated whether a similar phenotype is observed with the TPX2 phosphomutants. Compared to parental cells (Fig. 7C left & Fig. 7D), cells depleted of TPX2 have unstable kinetochore fibres (Fig. 7C right). Cells expressing either WT, or S738D, largely restore kinetochore fibre cold stability (Fig. 7C and D). In contrast, the S738A mutant has an impaired ability to rescue, with many S738A cells showing decreased kinetochore fibre stability. Taken together, we conclude from these data that phosphorylation of TPX2 at S738 is important for its mitotic functions.

Because phosphorylation of TPX2 at S738 is observed to be increased in the G2-phase fraction, we next investigated whether TPX2 S738 phosphorylation has a function in earlier mitotic events. One of the earliest events of mitosis is centrosome separation, which occurs prior to nuclear envelope breakdown. Plk1 and CDK activities promote Eg5-driven centrosome separation during late G2 and prophase (Smith et al. 2011). WT TPX2, which is nuclear in interphase (Trieselmann 2003), partially localises to centrosomes and microtubules in prophase (Ma et al. 2011). To assess whether TPX2 phosphorylation may play a role in prophase centrosome dynamics, we measured centrosome separation normalized to nuclear area (Fig. 8A) at nuclear envelope breakdown in cells overexpressing either WT, S738D, or S738A TPX2 (Fig. 8 B & C). Interestingly, overexpression of the TPX2 S738A mutant significantly decreases centrosome separation distance, compared either to parental cells, or to cells overexpressing either WT, or S738D TPX2 (Fig. 8B). In addition to the centrosome separation defect, spindle phenotypes manifest after nuclear envelope breakdown in cells overexpressing the TPX2 S738A mutant, as compared to cells overexpressing either TPX2 WT, or the S738D mutant (Fig. 8D & E). Cells overexpressing the TPX2 S738A mutant notably show a monopolar spindle phenotype (Fig. 8D bottom). The phenotype penetrance, i.e. the ratio of cells showing monopole vs. bipole spindles, increases with GFP fluorescence bin (Fig. 8F), suggesting that the phenotype is associated with high levels of expression of the TPX2 S738A mutant protein.

**Figure 8.**
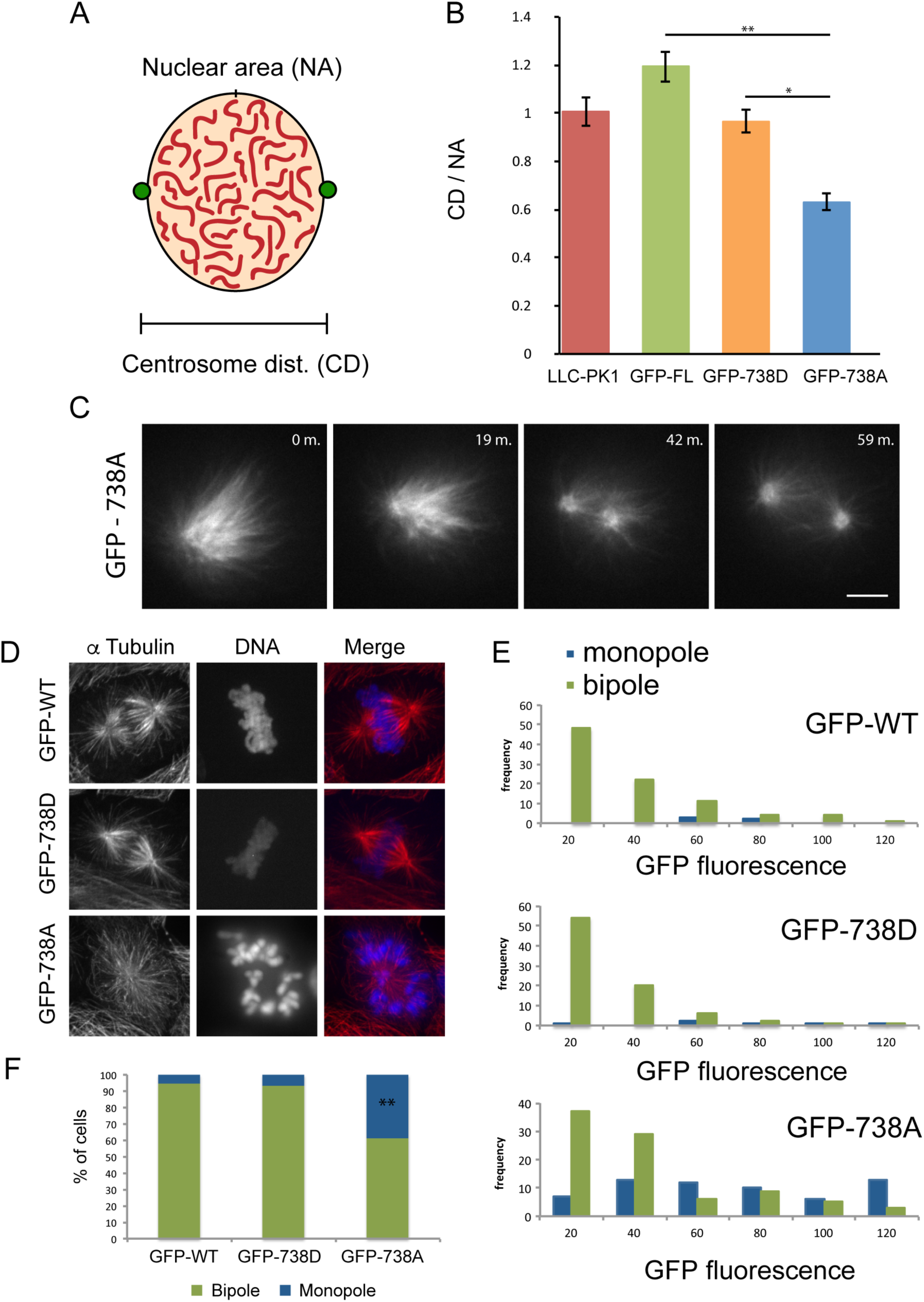
Overexpression of S738A impairs centrosome separation and formation of bipolar spindles. A) Schematic diagram illustrating the parameters measured. B) Measurement of centrosome separation in prophase cells expressing the indicated construct. N = 20, 6, 3, and 18, respectively. C) Live cell imaging of LLC-Pk1 cell overexpressing GFP-TPX2-S738A. Initially a monopolar spindle forms that subsequently forms a bipole. GFP fluorescence. D) Spindle phenotype in cells transfected with full-length wild-type GFP-TPX2, GFP-TPX2-S738A or GFP-TPX2-S738D. E) Quantification of overexpression phenotype shown in (D). Histograms showing frequency of monopoles and bipoles binned by level of GFP fluorescence (X-axis); green - bipole, blue – monopole. F) Bar graph showing all cells scored in B; expression of TPX2-S738A results in a 4-fold increase in monopolar spindles. N = 90, 95 and 150, respectively. D) Scale bar = 5 *μ* m. Time in minutes in upper right. **, P <0.001. * P < 0.05.

In summary, we conclude that high levels of a TPX2 mutant that cannot be phosphorylated on residue 738 significantly impairs centrosome separation, an early mitotic event.

### PRIMMUS analysis of protein abundance variation during mitosis

We next used the PRIMMUS method to perform a proteomic analysis of protein abundance variation across four temporally distinct stages of mitosis, reflecting prophase, prometaphase (1 & 2, see below) and anaphase. We aimed to screen for proteins that show abundance patterns resembling cyclins A and B, which could reveal novel targets whose degradation is also regulated during mitosis. We chose H3S28ph and cyclin A (CycA), as two markers with which to distinguish different mitotic subphases. Like H3S10ph, the H3S28ph signal is associated with chromatin condensation (as reviewed in (Hans & Dimitrov 2001)). During mitosis, cells undergo reversible condensation of chromatin, with highest levels of compaction observed during prometaphase and metaphase (Hans & Dimitrov 2001). Thus, cells showing the highest levels of H3S28ph signal (H3S28ph-high), represent prometaphase and metaphase cells, while cells showing intermediate levels of H3S28ph signal (H3S28ph-mid), are in early (prophase) and late (anaphase and telophase), stages of mitosis, respectively. Meanwhile, CycA is targeted for degradation by the APC/C during prometaphase in a SAC-independent manner (Elzen & Pines 2001). Thus, comparing either the presence, or absence, of CycA provides a means for distinguishing between ‘early’ (prometaphase and before) vs. ‘late’ (prometaphase and after) mitotic cells, respectively. Consistent with this, flow cytometry analysis of cells co-immunostained for H3S28ph and CycA show four subpopulations (labelled P1 – P4), which are H3S28ph positive (Fig. 9A).

**Figure 9.**
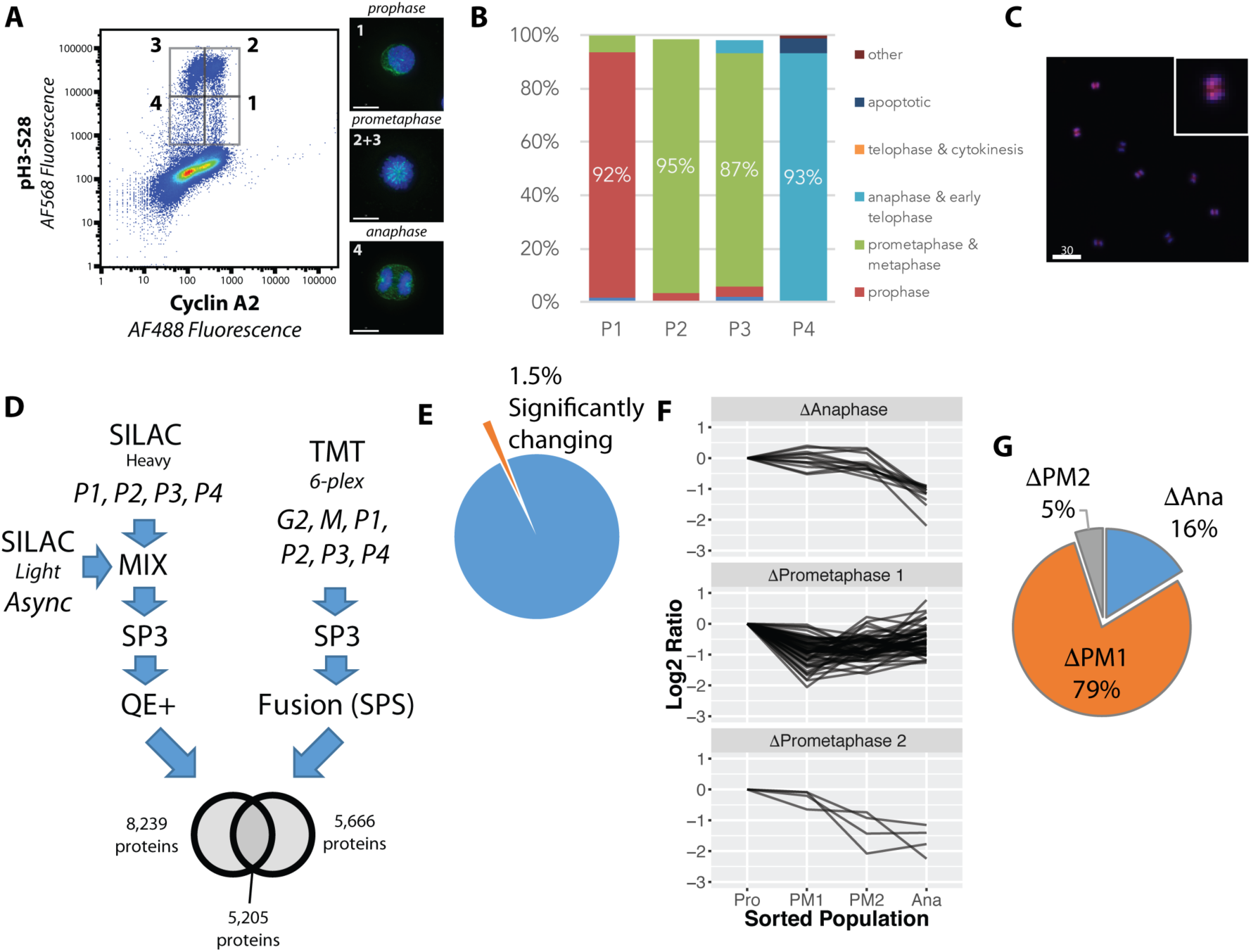
Proteome-wide analysis of protein abundance changes between mitotic subphases. A) Flow cytometry analysis of NB4 cells immunostained for pH3-S28 and CycA. Gates show populations collected by FACS. A, right) Representative light microscopy images of cell fractions. B) The frequency of each intra-mitotic stage was counted and quantified with 100 cells or more. C) Wider field of view of population 4, Ie., the anaphase-enriched population. D) Workflow for MS-based proteomic analysis involving SILAC and TMT based labelling and 3 biological replicates, resulting in 8,700 proteins identified in total. E) Pie chart indicating the percentage of significantly changing proteins (see text for criteria for significance). F) K-means clustering of profiles were qualitatively agglomerated into three groups based on subpopulation where ‘trough’ in abundance occurs. G) A pie chart showing the number and proportion of proteins in each protein group indicated in (F).

The four subpopulations described above were isolated by FACS and analysed by immunofluorescence microscopy (representative images are shown in Fig. 9A, right). The frequency of interphase, prophase, prometaphase, anaphase and telophase cells were each measured, with at least 100 cells counted for each subpopulation (Fig. 9B). This confirms that high enrichment efficiencies were obtained for prophase, prometaphase and anaphase, respectively. Telophase and cytokinesis cells were not observed in these subpopulations. We note that the gating strategy employed, which removes potential doublets from the analysis, biases against these cells (unpublished observations). Fig. 9C shows a representative image of the P4 subpopulation, showing high enrichment of anaphase cells. Based on these high enrichment efficiencies, we have relabeled these populations according to the major enriched phase represented, i.e., prophase (Pro), prometaphase 1 (PM1), prometaphase 2 (PM2) and anaphase (Ana), respectively.

The FACS protocol was repeated, using three separate asynchronous cell populations cultured and harvested on different days. Two of these populations were metabolically-labelled with stable isotope labelled amino acids for SILAC-based relative quantitation. These populations were sorted into the four subpopulations, P1 – P4, as described above. A third population was cultured using regular culture media containing non-dialysed bovine serum. For this third experiment, two other subpopulations were collected, i.e., G2 cells (i.e., 4N DNA content and pH3 negative cells) and M-phase cells (4N DNA content and pH3 positive cells), in addition to the P1 – P4 populations. Relative quantitation was performed using 6-plex peptide TMT labeling (Thompson et al. 2003). To maximize peptide recovery for MS analysis, the SP3 paramagnetic bead cleanup method was used (Hughes et al. 2014). SILAC samples were analysed using a Q-Exactive Plus, as previously described (Endo et al. 2017). TMT samples were analysed using a Fusion Tribrid instrument, using synchronous precursor selection (SPS) (McAlister et al. 2014) to minimize ratio compression. Thus, two distinct relative quantitation strategies were employed to reduce the likelihood of any potential artefacts from a single quantitation strategy to systematically bias the dataset.

Quantitative data for >8,700 proteins were obtained in the combined dataset across three biological replicates. 5,205 proteins (60%) were detected in both the TMT and SILAC datasets. To identify proteins whose abundance changed significantly, protein profiles were calculated based on the mean of the three biological replicates and ratios calculated, relative to prophase levels. Proteins showing missing values in any of the four mitotic subpopulations were discarded, leaving 5,340 proteins. A maximum fold change was calculated and cutoffs were established using Z-scores (∼1.8-fold at 95% confidence), which identified 235 proteins meeting this cutoff. Positive correlation between any two biological replicates was used as a second criterion for significance, with 136 proteins meeting these stringent criteria (Fig. 9E). Several proteins were excluded from the analysis due to missing values, which is a known technical issue with data-dependent acquisition. However, an alternative, non-mutually exclusive explanation, is the lack of detection reflects that the levels of these proteins fall below the detection limit due to physiological down-regulation. We reasoned that the latter explanation is more likely when proteins show missing values reproducibly in a single fraction. Two proteins met this criterion and were added to the candidate list (Supplementary Table X), which now totals 138 proteins. However, these two proteins were excluded in further clustering analysis due to missing values being incompatible with the standard k-means clustering algorithm. The vast majority of proteins (n = 5,202, ∼98.5%), showed no significant change (Fig. 9E).

The mean profiles for the 136 proteins showing the most significantly changing abundance levels were clustered using k-means, with the number of initial clusters (12) determined using a within-cluster sum-of-squares analysis. Within this dataset we were interested in identifying candidates for targeted protein degradation during mitosis. We therefore focused on those clusters where a decrease in protein abundance was evident. These were manually agglomerated into three clusters, based on the phase in which the decrease is observed (Fig. 9F). Most of these proteins (79%), show a decrease in abundance coincident with the CycA+ prometaphase subpopulation (PM1), with fewer proteins (16%), decreasing in anaphase and fewer still (5%) decreasing in the CycA-prometaphase subpopulation (PM2).

We next examined the mean protein abundance profiles for cyclins A and B (Fig. 10A, s.d. shown as a gray ribbon). As expected from the flow cytometry analysis and the sorting strategy employed, cyclin A2 shows high levels in Pro and PM1 and a marked decrease in PM2 and Ana. Two isoforms of cyclin B are detected (B1 and B2). The abundances of both isoforms remain relatively constant between Pro, PM1 and PM2 and decrease in Ana to 25-35% of prophase levels. These data are consistent with the targeting of cyclin B for degradation by the APC/C at the metaphase to anaphase transition. In contrast, the abundance of GAPDH, a protein that is not expected to be targeted for degradation during mitosis, is unchanged between these subpopulations.

**Figure 10.**
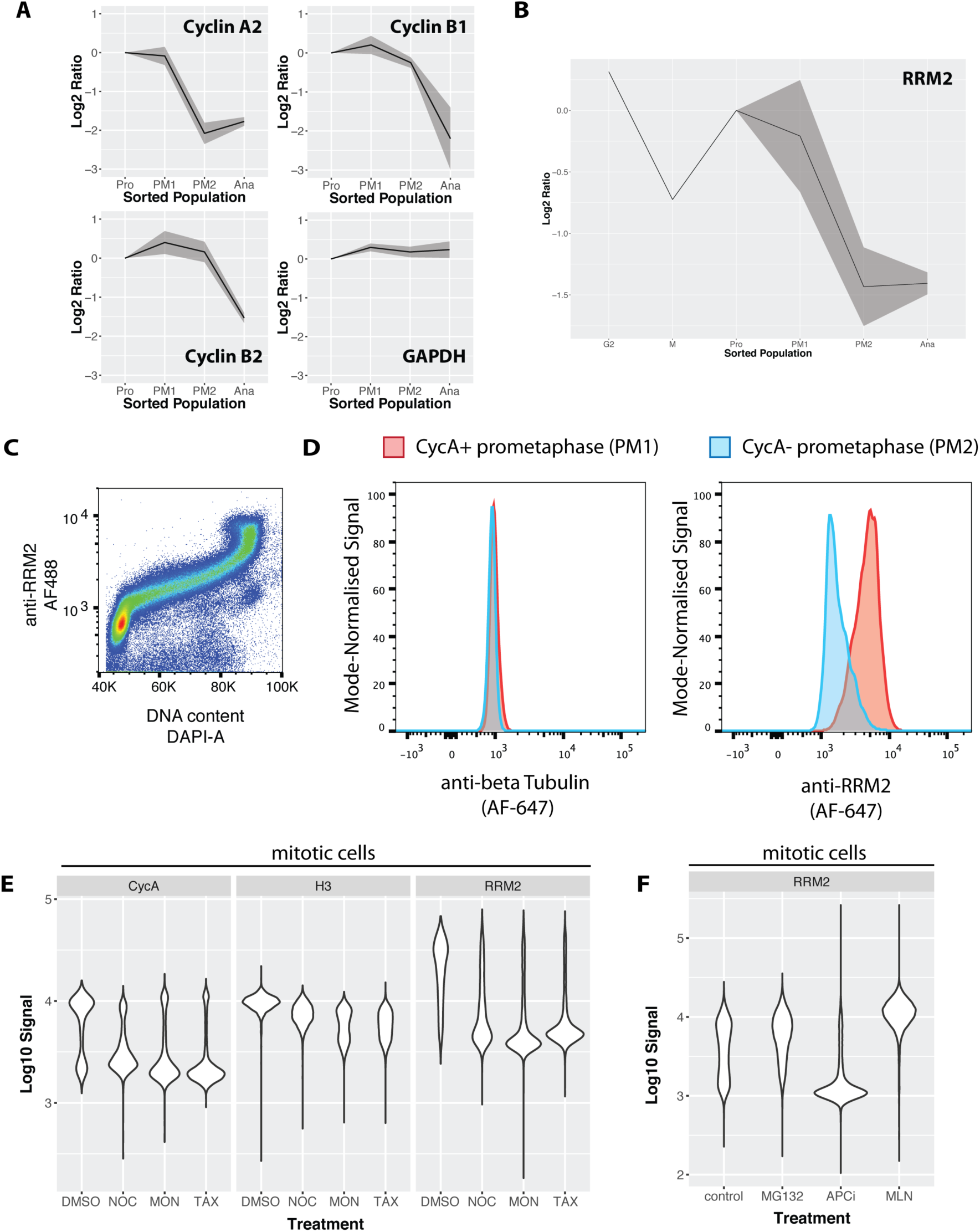
Ribonucleotide reductase M2 (RRM2) is degraded during prometaphase in a proteasome-dependent, MLN-4924 sensitive manner. A) Line graphs showing mean abundance profiles for Cyclin A2 (CycA), Cyclin B1, Cyclin B2, and GAPDH. Grey ribbons indicate 1 standard deviation from the mean. B) Mean abundance profile for RRM2. C) Flow cytometry analysis RRM2 levels vs. DNA content. D) Flow cytometry-based comparison of beta-tubulin (negative control, left) and RRM2 (right) levels in CycA+ (red) vs. CycA- (blue) prometaphase cells. E) Violin plots showing CycA, histone H3, and RRM2 levels in cells treated with either DMSO or microtubule drugs that activate the spindle assembly checkpoint (nocodazole, monastrol, taxol). F) Violin plots showing levels of RRM2 in cells treated with either vehicle control (DMSO), MG132, apcin+proTAME, or MLN4924.

### RRM2 is degraded during prometaphase via a MLN-4924-sensitive proteasomal pathway

We were interested in examining other proteins that co-clustered with the cyclin proteins. In the literature, there are few examples of substrates targeted for degradation during prometaphase, as compared with anaphase. We are aware of only two proteins, cyclin A2 and Nek2, for which there is significant evidence in the literature for targeted degradation during prometaphase(van Zon & Wolthuis 2010). Our data show 3 additional proteins clustering together with cyclin A2, i.e., ATAD2, GMNN and RRM2. Inspection of the protein abundance profiles show that while GMNN levels decrease during prometaphase to ∼60% of prophase levels, a second major decrease in GMNN abundance occurs during anaphase, where its levels drop to ∼20%. ATAD2, a protein involved in transcriptional co-activation of cell cycle genes, such as c-myc, cyclin D1 and E2F1, shows ∼50% reduction during prometaphase. ATAD2 contains a conserved, canonical D-box motif situated in a disordered linker region between a predicted DNA-binding bromo-domain and a second globular domain. ATAD2 was also recently shown to be cell cycle regulated via co-immunostaining of FUCCI-expressing cells with a rabbit polyclonal antibody recognizing ATAD2 (Human Protein Atlas, www.proteinatlas.org).

We investigated further the observed degradation of RRM2. RRM2 is an essential (Wang et al. 2015), cell cycle regulated subunit of ribonucleotide reductase that regulates the cellular deoxynucleotide pool (Nordlund & Reichard 2006). Properly timed degradation of RRM2 has been suggested to be important, because disrupting the normal degradation timing for RRM2 leads to genomic instability (D’Angiolella et al. 2012). RRM2 has been shown to be targeted for degradation by the APC/C-Cdh1 (Chabes et al. 2003) and Cyclin F/SCF (D’Angiolella et al. 2012) complexes during anaphase and in G1 and G2, respectively. Therefore, we were surprised to observe significant decreases in RRM2 abundance specifically during prometaphase, coincident with degradation of cyclin A2 (Fig. 10B).

A rabbit polyclonal antibody raised against RRM2 was validated by siRNA depletion (Fig. 10 Fig. Suppl. 1) and used to measure the levels of RRM2 across the cell cycle in NB4 cells. Consistent with its degradation by APC/C-Cdh1, flow cytometry analysis of RRM2 levels and DNA content show that RRM2 levels are low during G1 phase and increase significantly during S-phase, reaching a peak in G2&M (Fig. 10C). Interestingly, this analysis shows no significant decrease in RRM2 levels in G2, as was previously reported in HeLa cells (D’Angiolella et al. 2012). Co-immunostaining cells to detect both cyclin A and H3S28ph enables a flow cytometry-based comparison of RRM2 levels, using the same gating strategy as the MS-based analysis used to define subpopulations. A separate control sample, using an antibody recognizing alpha tubulin, was included as a negative control. Fig. 10D shows the fluorescence signal distributions for two subpopulations marked by H3S28ph and CycA staining: i.e., CycA+ prometaphase (PM1) versus CycA-prometaphase (PM2) cells. Comparing protein levels in the respective PM1 vs PM2 subpopulations, alpha tubulin shows no change, as expected (Fig. 10D, left). In contrast, the level of RRM2 in PM2 is markedly decreased, as compared with PM1, consistent with the MS-based quantitation (Fig. 10D, right).

We next explored potential mechanisms leading to targeted degradation of RRM2. Like CycA, the prometaphase decrease in RRM2 levels occurs in the presence of an active spindle assembly checkpoint (SAC) (Fig. 10E). Mitotic degradation of RRM2 is sensitive to 4 hrs of MG-132 treatment, suggesting that the degradation occurs via the proteasome pathway. Inhibition of APC/C activity, using combined treatment with apcin and proTAME (Sackton et al. 2014), was sufficient to block CycB degradation, but insufficient to stop CycA and RRM2 degradation in NB4 cells. In contrast, treatment with MLN-4924 (Soucy et al. 2009), an inhibitor of NEDDylation and cullin ring ligases (CRLs), completely blocks the decrease of RRM2 levels in prometaphase and leads to levels either similar, or even slightly higher, than observed in control cells (Fig. 10F). As expected, the decrease in CycA in prometaphase is largely unaffected in MLN-4924 treated cells (data not shown).

We conclude that RRM2 is targeted for proteasomal degradation during prometaphase via a MLN-4924-sensitive pathway, probably involving an SCF complex. As discussed further below, a likely candidate is the cyclin F/SCF complex, as RRM2 has been described previously as a substrate of this E3 ligase (D’Angiolella et al. 2012).

To maximize community access to the entire dataset described in this study, including the optimization experiments and the measurements of protein accumulation across the cell cycle, all of the proteomics data are freely available via several outlets, including the Encyclopedia of Proteome Dynamics (http://www.peptracker.com/epd/) (Fig. 11). The EPD provides a searchable, open access database containing also proteome measurements from multiple large-scale experiments on human cells and model organisms (Larance et al. 2013). The data are also available at various stages of analysis, including raw MS files and MaxQuant-generated output (ProteomeXchange) and analysed data (Supplementary Tables).

**Figure 11.**
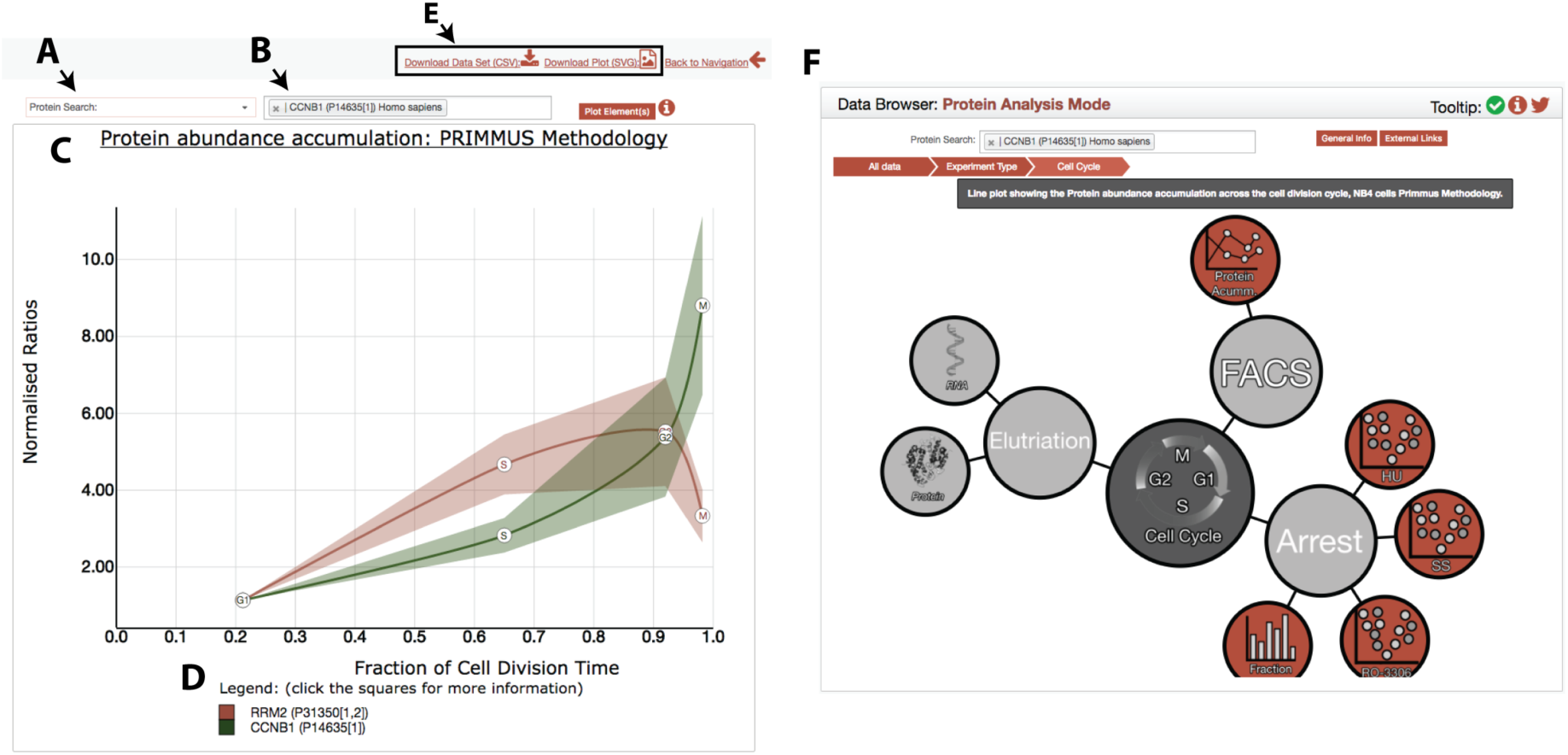
The PRIMMUS cell cycle data is accessible through the Encyclopaedia of Proteome Dynamics (EPD). User interface features include: A) specifying type of search, including search for individual proteins, GO term, and CORUM complex membership, B) an input box for protein and other (e.g. GO term) identifiers, C) a ribbon graph showing plots for input protein identifiers (here for illustration are shown RRM2 and CCNB1) with lines indicating mean profile and ribbons indicating s.e.m., D) an interactive legend to show more information on individual proteins, and E) options to output visualisation as an SVG file or underlying data in CSV format. F) Data from other cell cycle proteomic datasets from the Lamond laboratory can be easily visualized for the same proteins using the navigation bubble map.

## Discussion

A major challenge with the biochemical analysis of mitotic cells is that each subphase of mitosis is exceedingly short, lasting only minutes or less (Sullivan & Morgan 2007). Additionally, there is significant cellular heterogeneity in phase dwell times. Timing heterogeneity, which can be accounted for in timelapse imaging studies (Akopyan et al. 2014), present major technical challenges to synchronization-based strategies for biochemical analysis. In this study, we characterize the PRIMMUS method, which provides a flexible approach that facilitates quantitative proteomic studies on specific cell subsets isolated by FACS on the basis of staining for *intracellular* antigens. For separation of interphase cells (G1, S, G2), centrifugal elutriation is an alternative to FACS, but provides lower resolution separation, is not applicable to all cell types and does not efficiently separate G2 and M phase cells. We therefore used FACS to produce highly enriched populations of cells at specific cell cycle stages. Cells growing in asynchronous cultures were FACS separated by either, a) DNA content and phosphorylation of histone H3, obtaining high purity populations of G1, S, G2 and M phase cells, or by b) DNA content, phosphorylation of histone H3 and the degradation of CycA, obtaining high purity populations of prophase, prometaphase and anaphase intra-mitotic cells. Using these isolated cell populations, we provide the first specific MS-based proteomic analysis of intra-mitotic phase cells isolated from asynchronously growing cultures.

We validated the PRIMMUS method by demonstrating that global MS-based protein identification and quantitation is compatible with the analysis of populations of fixed cells that have been permeabilised, stained to detect *intracellular* antigens and isolated by FACS. While FACS has been used previously in conjunction with RNA-seq to compare mRNA abundances of cell subsets (Hrvatin et al. 2014), this study provides the first example we know of where permeabilised, fixed and intracellular immunostained cells have been FACS sorted and used for quantitative, MS-based proteome analysis. In principle, the PRIMMUS approach can be used to characterize any distinct type of cell subpopulation that can be defined using one or more diagnostic antigens, an abundance differential for a specific epitope, or combination of epitopes, including intracellular and intranuclear antigens. We also show that PRIMMUS enhances the sensitivity of quantitative proteomics technology to detect either changes in abundance, and/or changes in other protein properties, such as post-translational modifications, because it facilitates the analysis of the specific subsets of cells in which the change occurs, without diluting this signal by analyzing mixed populations, including non-responding cells. This is illustrated here by our demonstration of up to a five-fold sensitivity gain in detecting cell cycle regulated protein abundance changes, as judged by comparing data obtained using PRIMMUS with centrifugal elutriation.

We have recently shown by proteomic analysis of NB4 cells after arrest at specific cell cycle stages by drug treatments that the stress resulting from drug arrest causes major changes in the proteome distinct from the physiological regulation of protein levels during unperturbed cell cycle progression (Ly et al. 2015). In the PRIMMUS method, fixation by FA captures cells ‘frozen in motion’ and thereby prevents significant perturbation of cellular physiology due to extended sample handling. Consistent with this idea, we previously showed that that the abundance of the apoptosis-associated protein BID is upregulated by the CDK1 inhibitor RO-3306, which arrests cells at the G2 and M border (Ly et al. 2015). In this study, using PRIMMUS, we observe BID levels stay relatively constant across the unperturbed cell cycle, consistent with our results comparing minimally perturbed cells separated by centrifugal elutriation (Ly et al. 2014). While FACS can also be used to sort live cells, thereby avoiding procedures involving fixation and permeabilisation, the process of FACS-based separation of live cells can induce cellular stress and this in turn can rapidly alter the proteome. The proteomes of live cells immediately after FACS may thus be significantly remodeled by activation of cellular stress responses. Furthermore, most live cell strategies require the construction of transformed cell lines expressing one or more fluorescent-tagged fusion proteins. In contrast, the PRIMMUS approach described here avoids any requirement for the use of exogenous, tagged proteins and provides a general strategy that can be applied to both cultured cell lines and primary cells isolated *ex vivo*.

We used PRIMMUS, combined with high accuracy cell mixing and quantitative metabolic labeling, to measure and compare the abundances of thousands of individual proteins across an unperturbed cell division cycle, including different substages of mitosis. Our data show that while exponential accumulation is observed for bulk protein, the rate of accumulation of individual proteins can deviate significantly from an exponential growth fit. For example, histones, which constitute ∼3-5% of the total protein abundance in NB4 cells (Ly et al. 2014), deviate from bulk protein accumulation by remaining flat between G2 and M, consistent with the synchronization of DNA replication and histone synthesis (Nelson et al. 2002),(Robbins & Borun 1967). Intriguingly, for p53 and some other proteins associated with stress responses and/or apoptosis, protein abundance remains relatively flat across the cell division cycle. Due to cell cycle-dependent increases in cell size and bulk protein abundance, the effective net intracellular concentrations of these proteins must thus decrease as the cell cycle progresses. Any functional consequence of a decrease in their abundance may however be offset if mechanisms exist to create local concentration hotspots at specific subcellular locations.

Detailed proteomic analysis of protein abundances across prophase, prometaphase and anaphase cells showed that only ∼1.5% of the proteins quantitated significantly decrease in abundance. These proteomic data showed RRM2 levels decreasing during prometaphase. Consistent with this, subsequent flow cytometry analysis showed that RRM2 and CycA levels are correlated during mitosis. The decrease in RRM2 levels was prevented by treatment of cells with either MLN-4924, or MG-132, consistent with the mechanism causing a decrease in RRM2 levels involving a cullin E3 ubiquitin ligase and proteasomal degradation. It has been shown that RRM2 is targeted for degradation by Cyclin F/SCF during G2 (D’Angiolella et al. 2012) and several studies have shown co-immunoprecipitation of RRM2 and Cyclin F in asynchronous cell extracts (D’Angiolella et al. 2012; Huttlin et al. n.d.; Hein et al. 2015). It has also been reported that disruption of the Cy motif in RRM2, which is important for recognition by Cyclin F/SCF, stabilizes RRM2 levels and increases DNA pools and genome mutation frequencies (D’Angiolella et al. 2012). It will be interesting in the future to investigate further the timing of RRM2 degradation and its relation to maintaining genome stability.

We show that PRIMMUS is compatible also with analysis of cell cycle regulated protein phosphorylation by comparing phosphorylation in G1-, S-, G2- and M-phase enriched fractions purified by FACS. While essentially all significantly changing phosphorylation sites peak during mitosis, we identify a subset of phosphorylation sites that show increased phosphorylation in the G2-enriched fraction, which we call ‘early risers’. These early risers share functional similarities, including enrichment in the ‘classic’ CDK substrate phosphorylation motif and localization to chromatin and the nuclear envelope. We suggest that these early rising sites may be the downstream effect of mitotic entry kinase activity, which includes Plk1 and Cdk1 (Smith et al. 2011). Interestingly, the sites that change by the highest amount in G2, besides factors strongly associated with DNA replication, are localized to the nuclear pore and nuclear envelope. Additionally, several sites, such as on lamina-associated polypeptide 2 (LAP2) and NUP153, contain both Plk1 and Cdk substrate sequence motifs (Supplementary Table 2). An expanded analysis of phosphorylation during S- and G2-phases will be an important goal of future studies and will help to define the targets of the mitotic entry network in detail. This may provide clues to the sequence of signaling events that precede nuclear envelope breakdown.

When the phosphoproteomic dataset from this study was compared with a previously reported atlas of mitotic phosphorylation in human cells derived using nocodazole arrested HeLa cells (Olsen et al. 2010), some phosphorylation sites were identified that changed in one dataset, but not the other. The sites that differed between the two datasets were preferentially on proteins with specific functional GO annotations, including ‘cell cycle’ and ‘mitosis’ for phosphorylation sites that are changing exclusively in the mitotic fraction in this PRIMMUS dataset, but instead proteins with phosphorylation sites that are changing exclusively in the mitotic fraction in the previously reported HeLa dataset are enriched for the annotation ‘splicing’ and not ‘cell cycle’ or ‘mitosis’. Several differences between these studies could explain study-specific phosphorylation changes. For example, cell-type specificity in cell cycle regulated protein phosphorylation, e.g. NB4 cells (this study), as compared with HeLa cells used in the previous study. An alternative, non-mutually exclusive, explanation could be the difference in the methods used for obtaining mitotic cells in the respective studies, i.e. FACS (this study) vs nocodazole-arrest (HeLa dataset (Olsen et al. 2010)). Previous work has shown that cell cycle arrests can induce proteomic changes that are not observed in a minimally perturbed cell cycle (Ly et al. 2015). Further analysis will be required to investigate the basis for the observed study-specific phosphorylation changes.

We show that for the TPX2 protein, the phosphorylation increase observed during G2 and mitosis has functional consequences on protein function and proper mitotic progression. Thus, mutation of the early rising phosphorylation site, S738, on TPX2 to a non-phosphorylatable residue (alanine), elicits a significant reduction in centrosome separation and defects in bipolar spindle assembly, when overexpressed. The C-terminal 37 amino acids (including S738) have been shown to be important in recruiting Eg5 to spindle microtubules and modulating Eg5-dependent microtubule gliding activity (Ma et al. 2011). Replacement of endogenous TPX2 with the S738A mutant shows phenotypes intermediate between wild-type and a TPX2 C-terminal deletion mutant lacking the Eg5 interaction domain. Interestingly, both Eg5 and the TPX2 S738A mutant correctly localize to spindle poles and spindle microtubules during mitosis (unpublished observations), suggesting that phosphorylation does not prevent TPX2 from recruiting Eg5 to spindle microtubules. However, the S738A phenotypes suggest that TPX2 S738 phosphorylation plays a key role in affecting the ability of TPX2 to modulate Eg5 activity.

In addition to the established functions of TPX2 in spindle assembly during prometaphase, evidence in plant cells suggest a second role during early prophase (Vos et al. 2008). In these cells, microinjection of anti-TPX2 antibodies during prophase inhibits nuclear envelope breakdown. In mammalian cells, a subset of TPX2 localises to centrosomes during prophase (Ma et al. 2011). Our present data showing centrosome separation delays with a phosphodefective TPX2 mutant supports dual roles of TPX2, before and after nuclear envelope breakdown.

In addition to PTM analyses, the PRIMMUS workflow can be extended in future by combining it with other complementary approaches to extend the depth of proteomic analysis of the selected cell subpopulations beyond only abundance measurements, e.g. using MS-based techniques for identifying protein interaction partners and protein complexes (Kirkwood et al. 2013),(Kristensen et al. 2012). PRIMMUS is highly complementary to recent studies dissecting, in high resolution, the timing of mitotic kinase activities and quiescence control by automated fluorescence imaging of fixed, immunostained cells (Akopyan et al. 2014; Spencer et al. 2013). As FA fixed cells are compatible with downstream RNA-seq analysis and ChIP studies, combining proteomics on sorted cells also with these high-throughput approaches would expand the in-depth analysis of gene expression (transcriptome and proteome) of rare populations and/or biochemically well-defined subpopulations, such as specific hematopoeitic cell populations that cannot be separated on the basis of cell surface markers alone.

## Online methods

### Cell culture

The NB4 cell line was originally established from acute myeloid leukemia blast cells grown on bone-marrow stromal fibroblasts (Lanotte et al. 1991). NB4 cells were obtained from the Hay laboratory (University of Dundee). Cells were cultured at 37 °C in the presence of 5% CO2 as a suspension in RPMI-1640 (Life Technologies) supplemented with 2 mM L-glutamine, 10% v/v foetal bovine serum (FBS, Life Technologies), 100 units/ml penicillin and 100 μg/ml streptomycin (100X stock, Life Technologies). Cell cultures were maintained at densities between 1 × 10^5^ to 1 × 10^6^ cells/ml. For SILAC labeling, culture media without arginine or lysine (DCP) was supplemented with either ‘light’ (Arg0, Lys0, Cambridge Isotope Labs) or ‘heavy’ (Arg10, Lys8, Cambridge Isotope Labs) isotopomers of arginine and lysine. SILAC labeling media was additionally supplemented with dialysed serum (10% final), 1X insulin-transferrin-selenium (100X stock from Life Technologies), 1x MEM vitamins (100X stock from Life Technologies), and 90 mg/L proline.

### Mammalian depletion and rescue experiments

LLC-Pk1 cells were co-nucleofected using an Amaxa nucleofector and Mirus nucleofection reagent with siRNA targeting endogenous pig TPX2 (5’ GAAUGGUACAGGAGGGCUU 3’) and with a plasmid expressing GFP tagged si-RNA resistant human TPX2, or human TPX2 with S738 mutated to either A or D. Additionally, this experiment was performed using LLC-Pk1 cells expressing localization and affinity purification (LAP) tagged mouse TPX2 from a bacterial artificial chromosome (BAC). For generation of mutants at S738, the BAC was modified using a modified LAP cassette (Poser et al. 2008). BAC DNA was nucleofected into LLC-Pk1 cells and selection was performed as previously described (Ma et al., 2011). Prior to use in experiments, the cells were treated with siRNA to deplete endogenous pig TPX2. To overexpress TPX2 constructs, 2ug of GFP-TPX2-wild type or 738A or 738D DNA was nucleofected into LLC-Pk1 cells. Overexpressing cells were maintained in cell culture without selection. To assay kinetochore fiber cold stability, cells were incubated in 5 uM MG132 for 1.5 hours to arrest cells in metaphase and then washed twice in ice cold PBS, placed in ice cold culture medium for 10 minutes on ice. Cells were fixed and stained for tubulin.

### Immunofluorescence staining

NB4 cells (∼0.5 × 10^8^ cells) were washed once with phosphate-buffered saline (PBS) and resuspended in freshly prepared 50 ml 0.5% formaldehyde in PBS. Cells were fixed with formaldehyde for 30 min at room temperature with shaking. Cells were pelleted, and permeabilised with 50 ml cold 90% methanol. Cells were then stored at - 20°C prior to staining.

Fixed, permeabalised cells were washed once with PBS and resuspended in blocking buffer, 5% bovine serum albumin (BSA) in Tris-buffered saline (TBS) + 0.05% sodium azide. Cells were blocked for 10 min at room temperature, pelleted, and resuspended in primary antibody solution (1:200 in blocking buffer). Cells were stained with primary antibody overnight at 4°C. Stained cells were then washed twice with PBS, and stained with dye-conjugated secondary antibodies (1:200 in blocking buffer) for 1 hour at room temperature. Stained cells were washed twice with PBS, pelleted, and resuspended in DAPI solution (5 μg/ml in PBS).

LLC-Pk1 cells were rinsed twice in room temperature PBS and fixed in paraformaldehyde-glutaraldehyde fixative (3.7% paraformaldehyde, 0.1% glutaraldehyde) in PBS containing 0.5% Triton X-100; fixation was carried out for 10 min. Microtubules were stained using a mouse anti-tubulin antibody (DM1a) or rat anti-tubulin (YL ½) and appropriate secondary antibodies. Incubations with primary antibodies were performed for either 1 hour at 37 C or overnight at room temperature; secondary antibodies were used at room temperature for 45 min. Antibodies were mixed at the appropriate final dilution with PBS containing 2% BSA, 0.1% Tween and 0.02% sodium azide.

### Flow cytometric analysis of cell cycle distribution

Cells stained with either PI or DAPI were analysed on an LSR Fortessa flow cytometer and data acquired using DIVA software (Becton Dickinson). DNA content was evaluated based on DAPI fluorescence (measured using 355nm excitation and emission at 450±50nm) or PI fluorescence (measured using 488nm excitation and emission at 585±42nm). Doublet discrimination was used to remove cell doublets and clumps using DAPI/PI-A and DAPI/PI-W measurements. The cell cycle distribution of single (gated) cells was plotted as DAPI/PI-A. Data was analysed using Flowjo software (Treestar inc.) and cell cycle distributions determined using the Watson/Pragmatic model.

### Fluorescence Activated Cell Sorting (FACS)

Immediately prior to sorting, cells were passed through a 50μm filter to remove cell clumps. Cells were sorted on an Influx cell sorter (Becton Dickinson) using a 100μm nozzle and sheath pressure of 20psi. Cells were distinguished from particulate material on the basis of forward scatter and side scatter and single cells were identified on the basis of DAPI-W v DAPI-A measurements. During sorting, data was analysed using FACS Sortware software (Becton Dickinson).

### Immunofluorescence microscopy

Cells purified by FACS were settled onto poly-lysine coated coverslips (BD Biosciences) for 30 min at room temperature. The liquid was then carefully aspirated. Cells were fixed with 2% FA in PBS for 10 min at room temperature. Cells were washed twice with PBS and stained with primary antibodies for 1 hour at room temperature. Cells were washed twice with PBS and stained with dye-conjugated secondary antibodies for 30 min at room temperature. Cells were washed twice with PBS and stained with DAPI (5 μg/ml in PBS) for 1 min at room temperature. Cells were washed once with PBS and mounted in Vectashield medium (Vector Laboratories). Cells were visualized using a wide-field fluorescence microscope (Zeiss, Jena, Germany; Axiovert-DeltaVision Image Restoration; Applied Precision, LLC).

LLC-Pk1 cells were cultured in Hams F-10 mixed one to one with Opti-MEM, and containing 7.5% fetal calf serum. Prior to use in experiments cells were plated on 22X22 mm #1.5 coverslips in 35 mm dishes or on to the surface of glass bottom petri dishes (Mattek Corp). Cells were imaged using a Nikon A1R+ resonant scanning confocal system equipped with a 60X NA 1.4 objective lens. For live cell imaging, cells were imaged using a Nikon Eclipse TE300 or TiE equipped with a Yokagawa spinning disc scan head and 100X NA 1.4 objective lens as previously described(Ma et al. 2011). Image analysis was performed in ImageJ or FIJI.

### Cell lysis, reverse crosslinking, protein precipitation/SP3

Cells were resuspended in 4% SDS in PBS, homogenised with a probe sonicator (Branson, 10% power, 20 sec, 4°C), and heated to 95°C for 30 min to reverse crosslinks. For MS analysis, proteins were then reduced and alkylated using TCEP (25 mM final concentration, Sigma) and iodoacetamide (55 mM final concentration, Sigma). G1, S, G2, and M phase lysates for the single shot analyses were then chloroform-methanol precipitated. The mitotic substage fractions (G2, M, Pro, PM1, PM2, and Ana) were processed using the SP3 method, as described previously (Hughes et al. 2014). Phosphopeptide enrichment was performed using magnetic Ti:IMAC beads from Resyn Biosciences using the manufacturer’s protocol.

### Immunoblot analysis

Lysates for SDS-PAGE analysis were prepared in lithium dodecylsulphate sample buffer (Life Technologies) and 25 mM TCEP. Samples were heated to 65 °C for 5 min and then loaded onto a NuPage BisTris 4 – 12% gradient gel (Life Technologies), in either MOPS, or MES buffer. Proteins were electrophoresed and then wet transferred to nitrocellulose membranes at 35 V for 1.5 hours. Membranes were then blocked in 5% BSA in immunoblot wash buffer (TBS + 0.1% Tween-20) for 1 hour at room temperature. Membranes were then probed with primary antibody overnight at 4°C, washed and then re-probed with LiCor dye-conjugated secondary antibodies (either IRDye-688 or IRDye-800). Primary antibodies for cell cycle immunoblot analysis were obtained from Cell Signaling Technology (cyclin B1, cyclin A, cyclin E, CDT1). Bands were visualized using the Odyssey CLx scanner (LiCor Biosciences).

### LC-MS/MS analysis

Chloroform-methanol precipitated protein pellets were resuspended in 8 M urea in digest buffer (100 mM Tris pH 8.0, 1 mM CaCl_2_). The protein solution was then diluted to 4 M urea with digest buffer and then digested with Lys-C, which was added at a 1:50 w/w Lys-C:protein ratio from a 1 mg/ml stock in water (Wako Chemicals) overnight at 37°C. The lysates were then further diluted with digest buffer to 0.8 M urea and digested with trypsin, which was added at a 1:50 w/w trypsin:protein ratio from a 0.2 mg/ml stock in 50 mM acetic acid (Thermo Pierce) for four hours at 37°C. The digests were then desalted using SepPak-C18 SPE cartridges, dried, and resuspended in 5% formic acid. Peptide concentrations were determined using the amine-reactive, fluorogenic CBQCA assay (Life Technologies).

SP3-processed proteins were digested in a similar manner to above. For one biological replicate, the resulting peptides were TMT-labelled according to manufacturer’s instructions. Peptides (SILAC or TMT) were then mixed, dried, and resuspended in high pH reverse phase buffer A (10 mM ammonium formate in 2% ACN, pH 9.3). Peptides were then chromatographed on a Dionex Ultimate 3000 off-line HPLC equipped with a XBridge Peptide BEH C18 4.6 mm x 250 mm column packed with 3.5 μm particles containing 130 angstrom pores (Waters) and a standard gradient over 20 min from 25% at 0 min to 60% B at 11 min (10 mM ammonium formate in 80% ACN, pH 9.3) at a flow rate of 1 ml/min. 48 fractions were collected into 24 wells in a concatenated format.

SILAC-labelled peptides were analyzed using a Dionex RSLCnano HPLC-coupled Q-Exactive Orbitrap mass spectrometer (Thermo Fisher Scientific). Peptides were first loaded onto a 2 cm PepMap trap column in 2% acetonitrile + 0.1% formic acid. Trapped peptides were then separated on an analytical column (75 μm x 50 cm PepMap-C18 column) using the following mobile phases: 2% acetonitrile + 0.1% formic acid (Solvent A) and 80% acetonitrile + 0.1% formic acid (Solvent B). The linear gradient began with 5% B to 35% B over 220 min with a constant flow rate of 200 nl/min. The peptide eluent flowed into a nanoelectrospray emitter at the front end of a Q-Exactive Plus (quadrupole Orbitrap) mass spectrometer (Thermo Fisher Scientific). A typical ‘Top10’ acquisition method was used. Briefly, the primary mass spectrometry scan (MS1) was performed in the Oribtrap at 70,000 resolution. Then, the top 10 most abundant m/z signals were chosen from the primary scan for collision-induced dissociation in the HCD cell and MS2 analysis in the Orbitrap at 17,500 resolution. Precursor ion charge state screening was enabled and all unassigned charge states, as well as singly charged species, were rejected.

TMT-labelled peptides were analyzed using a Dionex RSLCnano HPLC-coupled Tribrid Fusion mass spectrometer (Thermo Fisher Scientific). Peptides were first loaded onto a 2 cm PepMap trap column (100 μm) in 2% acetonitrile + 0.1% formic acid. Trapped peptides were then separated on an analytical column (75 μm x 50 cm PepMap-C18 column) using the following mobile phases: 2% acetonitrile + 0.1% formic acid (Solvent A) and 80% acetonitrile + 0.1% formic acid (Solvent B). The linear gradient began with 5% B to 35% B over 220 min with a constant flow rate of 200 nl/min. The peptide eluent flowed into a nanoelectrospray emitter at the front end of either a Q-Exactive, or Q-Exactive Plus (quadrupole Orbitrap) mass spectrometer (Thermo Fisher Scientific). A max cycle time acquisition method was used (2s). The primary mass spectrometry scan (MS1) was performed in the Oribtrap at 120,000 resolution. Then, the top N most abundant m/z signals were chosen from the primary scan for CID (30%) and Rapid-mode MS2 analysis in the linear ion trap. A synchronous precursor selection (SPS) method was employed. MS2 product ions were selected using 4 notches with a maximum injection time of 300 ms, fragmented by HCD (55%), and TMT tags mass analysed in the Orbitrap at 60,000 resolution. Precursor ion charge state screening was enabled and all unassigned charge states, as well as singly charged species, were rejected.

### MS data analysis

The SILAC and TMT RAW data files produced by the mass spectrometer were analysed using the quantitative proteomics software MaxQuant, versions 1.5.1.2 and 1.5.3.30 (Cox & Mann 2008). This version of MaxQuant includes an integrated search engine, Andromeda (Scheltema et al. 2011). The database supplied to the search engine for peptide identifications was a UniProt human protein database (‘Human Reference Proteome’ retrieved on April 16, 2016) combined with a commonly observed contaminants list. The initial mass tolerance was set to 7 p.p.m. and MS/MS mass tolerance was 20 ppm. The digestion enzyme was set to trypsin/P with up to 2 missed cleavages. Deamidation, oxidation of methionine and Gln->pyro-Glu were searched as variable modifications. Identification was set to a false discovery rate of 1%. To achieve reliable identifications, all proteins were accepted based on the criteria that the number of forward hits in the database was at least 100-fold higher than the number of reverse database hits, thus resulting in a false discovery rate of less than 1%. Protein isoforms and proteins that cannot be distinguished based on the peptides identified are grouped by MaxQuant and displayed on a single line with multiple UniProt identifiers. The label free quantitation (LFQ) algorithm in MaxQuant was used for protein quantitation. The algorithm has been previously described (Cox 2014). Protein quantitation was performed on unmodified peptides and peptides that have modifications that are known to occur during sample processing (pyro-Glu, deamidation). All resulting MS data were integrated and managed using PepTracker Data Manager, a laboratory information management system (LIMS) that is part of the PepTracker software platform (http://www.PepTracker.com).

## Acknowledgements

This work was supported by grants to AIL from the Wellcome Trust (105024/Z/14/Z, 108058/Z/15/Z) and the EU EpiGeneSys network (Grant#: HEALTH-F4-2010-257082). We thank the UK Research Partnership Investment Fund and the Scottish Funding Council (Project H13047) for proteomics instrumentation funding and the Wellcome Trust (097418/Z/11/Z) for supporting the Flow Cytometry and Cell Sorting Facility at the University of Dundee. We thank our colleagues in the Lamond group for advice and discussion. We thank Calum Thompson and Alan Prescott (Dundee Light Microscopy Facility), the Swedlow laboratory, and Raffaella Pippa (Lamond lab) for technical advice and assistance.

---

**Figure 1 Figure Supplement 1. Immunoblot analysis of the effect of FA on the electrophoretic migration of individual proteins.**

Immunoblot analysis of crosslinked lysates for alpha tubulin (A) and histone H3 (B). Immunoblot analysis of crosslinked & reverse crosslinked lysates for alpha tubulin (C) and histone H3 (D). Arrowheads indicate high MW bands remaining after reverse crosslinking step that migrate at a higher MW than the expected monomer mass (arrows).

**Figure 1 Figure Supplement 2. The effects of formaldehyde (FA) concentration on protein crosslinking and DNA staining.**

A) DNA content histograms from flow cytometry of cells fixed with the indicated concentrations of FA and using either propidium iodide (PI, left), or 4’,6-diamidino-2-phenylindole (DAPI, right), as the DNA-binding dye.

**Figure 10 Figure Supplement 1. siRNA-based validation of the anti-RRM2 antibody used in this study.**

## Supplementary Tables

**Supplementary Table 1. The effect of fixation and permeabilisation protocols on MS-based protein quantitation.**

A tab-delimited text file containing a list of all the protein groups identified with ≥2 peptides per protein group and their associated SILAC ratios comparing the different fixation and permeabilisation protocols.

**Supplementary Table 2. Analysis of protein accumulation across interphase and mitosis.**

The table consists of an excel file containing two worksheets. The first worksheet lists the protein groups identified with ≥2 peptides per protein group and their associated SILAC ratios in four biological replicates. For each biological replicate, the ratios were normalised to the ratio measured in G1. An offset was then added to the G1 ratio to account for the difference in time between cell division and an average G1 cell (calculated from Figure 5B). The second worksheet also lists the same protein groups, but with the unnormalised ratios. These are the ratios that were used to produce the ‘neeps’ plot in Figure 5A.

**Supplementary Table 3. Analysis of protein phosphorylation across interphase and mitosis.**

The table consists of a tab-delimited file containing the phosphorylation sites measured, quality measures (PEP, Score), and TMT ratios calculated relative to the G1 fraction from the two biological replicates. B – biological replicate, fc – fold change, repcor – Pearson’s correlation score between the ratio patterns of the two biological replicates

**Supplementary Table 4. Analysis of protein abundances during mitotic subphases.**

The table consists of a tab-delimited file containing the proteins identified, quality measures (Q-value, Score, number of peptides), TMT ratios calculated relative to the prophase fraction, and SILAC ratios calculated relative to the prophase fraction in biological duplicate. cor – Pearson’s correlation score between the ratio patterns of the three biological replicates (only mitotic subphases are compared). numcor – number of times the Pearson’s correlation score is greater than 0.

